# Metabolic basis for the evolution of a common pathogenic *Pseudomonas aeruginosa* variant

**DOI:** 10.1101/2022.01.14.476307

**Authors:** Dallas L. Mould, Mirjana Stevanovic, Alix Ashare, Daniel Schultz, Deborah A Hogan

## Abstract

Microbes frequently evolve in reproducible ways. Here, we show that differences in specific metabolic regulation explain the frequent presence of *lasR* loss-of-function mutations in the bacterial pathogen *Pseudomonas aeruginosa.* While LasR contributes to virulence, *lasR* mutants have been associated with more severe disease. A model based on the intrinsic growth kinetics for a wild type strain and its LasR^-^ derivative, in combination with an experimental evolution based genetic screen and further genetics analyses, indicated that differences in metabolism were sufficient to explain the rise of these common mutant types. The evolution of LasR^-^ lineages in laboratory and clinical isolates depended on activity of the two-component system CbrAB, which modulates substrate prioritization through the catabolite repression control pathway. LasR^-^ lineages frequently arise in cystic fibrosis lung infections and their detection correlates with disease severity. Our analysis of bronchoalveolar lavage fluid metabolomes identified compounds that negatively correlate with lung function, and we show that these compounds support enhanced growth of LasR^-^ cells in a CbrB-controlled manner. We propose that *in vivo* metabolomes are a major driver of pathogen evolution, which may influence the progression of disease and its treatment.

## Introduction

Quorum sensing (QS) is a mechanism of microbial communication that regulates the expression of a suite of genes in response to diffusible autoinducers in a population (Schuster & Greenberg, 2007; Schuster, Lostroh, Ogi, & Greenberg, 2003). Despite the importance of cell-cell communication for virulence (Rumbaugh et al., 2009) and high conservation across divergent phylogenies, key QS regulators in diverse species, such as *Pseudomonas aeruginosa*, *Vibrio cholerae*, and *Staphylococcus aureus*, frequently lose function (Dallas L. Mould & Hogan, 2021), due to recent missense and nonsense mutations, indels, or genome rearrangements. These paradoxical findings suggest that there may be connections between QS and other key physiological pathways that have yet to be revealed.

In *P. aeruginosa,* many isolates from humans, plants, and water sources have loss-of-function mutations in the gene encoding the transcription factor LasR (Groleau, Taillefer, Vincent, Constant, & Déziel; O’Connor, Zhao, & Diggle, 2021; Schuster et al., 2003), which is central to an interconnected QS network (Schuster et al., 2003). LasR^-^ isolates have been repeatedly observed in *P. aeruginosa* lung infections in people with cystic fibrosis (pwCF) (Smith et al., 2006), and LasR^-^ isolate detection is associated with more rapid lung function decline and more inflammation than in comparator populations (Hoffman et al., 2009; LaFayette et al., 2015). In a clinical study of acute corneal infections (Hammond et al., 2016), LasR^-^ strains also correlated with more damage and worse outcomes.

Multiple studies contribute to our understanding of the physiologies and social interactions that impact *lasR* loss-of-function mutant fitness. Several studies provide evidence in support of the model that LasR^-^ strains are “social cheaters” that reap the benefits of shared goods secreted by neighboring wild-type cells without incurring the metabolic costs (Sandoz, Mitzimberg, & Schuster, 2007). In this case, LasR^-^ strains grow better when the wild type is in the majority, and crash when a critical threshold of LasR^-^ cells is surpassed (West, Griffin, Gardner, & Diggle, 2006). The extent of *lasR* mutant “cheating” depends on the cost-benefit difference, and multiple shared goods, including siderophores, must be considered (Özkaya, Balbontín, Gordo, & Xavier, 2018). To combat the rise of cheaters, *P. aeruginosa* produces products such as hydrogen cyanide, rhamnolipids, or pyocyanin that inhibit growth of quorum sensing mutants through a process known as “policing” (Castañeda-Tamez et al., 2018; Rodolfo García-Contreras et al., 2020; M. Wang, Schaefer, Dandekar, & Greenberg, 2015).

There is evidence that the presence of LasR^-^ subpopulations may be beneficial (R. García-Contreras & Loarca, 2020) and lead to emergent properties including metabolite-driven interactions between wild type and *lasR* mutants that provoke the production of QS-controlled factors by the *lasR* mutant to levels greater than in wild-type monocultures (D. L. Mould, Botelho, & Hogan, 2020). In addition to the interactions between LasR^+^ and LasR^-^ cells that influence the fitness and behavior of LasR^-^ strains described above, there are important intrinsic characteristics of LasR^-^ strains including increased Anr-regulated microoxic fitness (Clay et al., 2020), resistance to alkaline pH in aerobic conditions (Heurlier et al., 2005), and altered metabolism (D’Argenio, Wu, Hoffman, Kulasekara, Déziel, et al., 2007). The metabolic advantages associated with LasR^-^ strains include growth on individual amino acids (D’Argenio, Wu, Hoffman, Kulasekara, Déziel, et al., 2007). The numerous differences described between LasR^+^ and LasR^-^ strains indicate that an understanding of the factors that drive the rise and persistence of *lasR* mutants may be complex and are not yet well understood.

Here, we use mathematical modeling, experimental evolution-based genetic screens, phenotype profiling, and whole-genome sequencing of evolved communities in different backgrounds to understand the rise of LasR^-^ strains over only a few serial passages. We identified the CbrAB pathway as the strongest contributor to the rise of *lasR* loss-of-function mutants, and our findings were not specific to strain background or medium. LasR^-^ strains are more commonly detected in samples from individuals with more severe CF lung disease (Smith et al., 2006). Analysis of bronchoalveolar lavage samples from pwCF and non-CF comparators identified several compounds that were higher in pwCF and that inversely correlated with lung function. LasR^-^ strains showed improved growth on the majority of these compounds, many of which were amino acids, and epistasis analysis confirmed that the improved growth was due to altered activity of the CbrB-CrcZ-Crc pathway.

## Results

### Mathematical model built from monoculture growth data predicts the observed rise of *lasR* loss-of-function mutants

Our previous work on microbial interactions involving LasR^+^ and LasR^-^ *P. aeruginosa* revealed subtle differences in growth kinetics (D. L. Mould et al., 2020). In monoculture, *P. aeruginosa* strain PA14 Δ*lasR* had no lag phase while the wild type had a lag phase of 1 h (**Fig. 1** for summary data and **Fig. S1** for growth curve). Furthermore, consistent with work by others (Diggle, Griffin, Campbell, & West, 2007; Rodolfo García-Contreras et al., 2020), the Δ*lasR* strain had a 16% lower growth rate but a 1.5-fold higher yield in LB. We found no differences in death rate resulting from elevated culture pH (as has previously been reported in low oxygen conditions (Heurlier et al., 2005)) or the onset of death phase relative to PA14 wild type under these conditions (**Fig. S1**).

**Figure 1.**
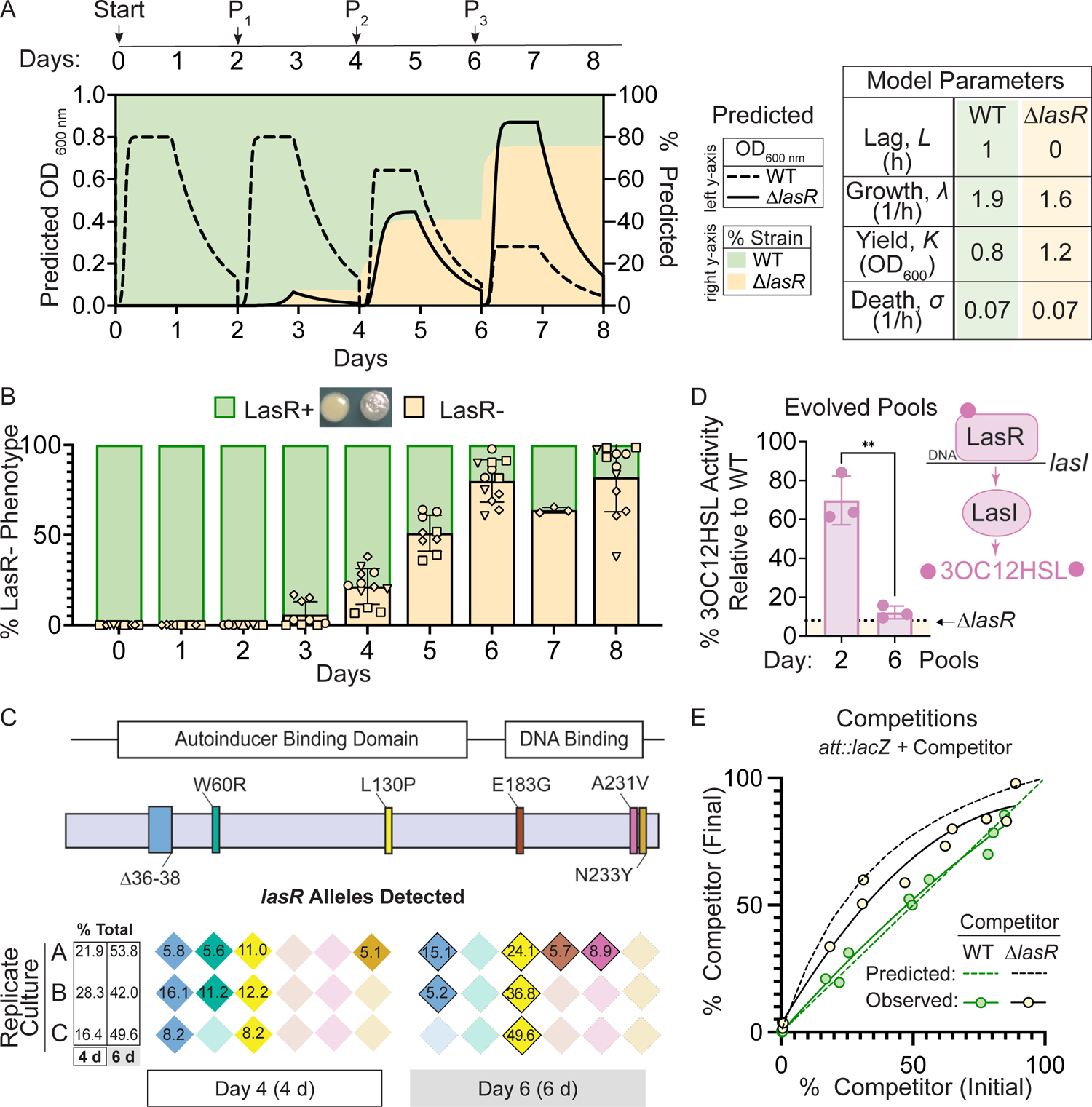
Mathematical model built from monoculture growth data is sufficient to explain the rise of LasR loss-of-function strains. A. Predicted densities (left y-axis) of mathematical model shown for wild type (WT, dashed line) and LasR-(solid line) strains. Predicted percentages (right y-axis) of LasR-(beige fill) and LasR+ (green fill) strains over the course of evolution regime in LB with passage (P_n_) every two days. Table shows experimentally-measured growth parameters obtained for strains PA14 WT and Δ*lasR* in LB used to create the model. B. Percentage of LasR-phenotypes observed in n 2: 4 independent evolution experiments in LB. A representative image of the smooth LasR+ and sheen LasR-colonies from Day 6 is shown. C. *lasR* alleles detected in the population at Day 4 (4 d) and Day 6 (6 d) by PoolSeq within the *lasR* coding sequence, which includes the autoinducer binding and DNA binding domains, for a representative experiment (diamond symbols, in B). The percentages of each allele and the sum indicated for each replicate culture. Each color represents a different allele. D. LasR regulates the production of its cognate autoinducer 30C12HSL via direct transcriptional control of the gene encoding the Lasl synthase. Lasl-produced autoinducer activity of evolved pools from a representative experiment (diamond symbols, in B) at days two and six. Activity is presented as a percentage of that produced by unevolved WT monocultures. The levels produced by the engineered Δ*lasR* control strain is shown for reference (dotted line). E. Comparison of predicted (dashed line) and observed (solid line) outcomes of competition assays initated at different initial ratios for which a constitutively tagged WT (*att*::*lacZ*) was competed against Δ*lasR* (beige, black line) or WT (control, green) competitors for 6 h (final) in planktonic LB cultures.

To determine if inherent differences in growth kinetics were sufficient to explain the rise of spontaneous LasR^-^ lineages, we built a mathematical model of strain competition based exclusively on experimentally-determined mono-culture growth parameters that predicted the relative changes in wild type and LasR^-^ cells grown on a common pool of growth substrates (**Fig. 1A**). We modeled cell density (**Fig. 1A**-left y-axis) and the percentage of LasR^-^ cells (**Fig. 1A**-right y-axis) assuming a shared nutrient source and passage every 48 hours which is a regime used previously to study the selection for LasR^-^ cells (Heurlier et al., 2005). Based on the mutation frequency of *P. aeruginosa* strain PA14 (0.52 × 10^−3^ per genome per generation) (Dettman, Sztepanacz, & Kassen, 2016) and the size of the *lasR* gene (720 bp) relative to the genome (∼6 Mbp), we approximate 50 *lasR* alleles with nucleotide changes would be present in a dense culture (∼ 10^8^ cells), a fraction of which would lead to a LasR^-^ phenotype. With the assumption of two to twenty LasR^-^ cells per inoculum (t=0, ∼ 10^5^ cells), the model predicted that ∼20% of the population would consist of LasR^-^ cells by Day 4, with increased percentages of ∼40% and ∼80% by Days 6 and 8, respectively (**Fig. 1A**) (Heurlier et al., 2005). Only minor differences in percentages resulted from changes in the initial LasR^-^ population.

We compared the model output to experimental data gathered with the same evolution regime. A single PA14 wild-type colony was used to inoculate a 5 mL culture of LB, which was grown to saturation and then used to inoculate three 5 mL LB cultures which were then passaged independently. Results from all three replicates from four independent experiments are shown. The percent of cells with LasR loss-of-function phenotypes were enumerated by plating and determining the percent of colonies with the characteristic “sheen” colony morphology of LasR^-^ cells that results from accumulation of 4-hydroxy-2-heptylquinoline (HHQ) (**Fig. 1B**) (D’Argenio, Wu, Hoffman, Kulasekara, Déziel, et al., 2007). In all four independent experiments, the percentage of colonies with the LasR^-^ phenotype rose from undetectable levels to an average of ∼80% over the course of eight days (**Fig. 1B**). To validate the use of colony sheen as an indicator of the LasR^-^ genotype, we evaluated ≥ 90 isolates with the characteristic LasR^-^ colony morphology for other phenotypes associated with LasR loss-of-function: low production of proteases and autoinducers (3OC12HSL and C4HSL). Most of the predicted LasR^-^ isolates (∼90%) had phenotypes that mirrored those of the PA14 Δ*lasR* strain, and not wild type (**Fig. S2**). Consistent with other studies (Feltner et al., 2016), approximately 15% of the cells with other LasR^-^ phenotypes produced high levels of C4HSL even though 3OC12HSL production was low.

The percentage of LasR^-^ cells predicted by the model matched the frequency of *lasR* alleles in genome sequence data from pools of colonies obtained from Day 4 and 6 cultures of a representative experiment (diamond symbols in **Fig. 1B**). Across replicates, six non-synonymous mutations were identified in *lasR* in the regions corresponding to LasR autoinducer binding (Δ36-38, W60R, and L130P) and DNA binding domains (E183G, A231V, and N233Y) (**Fig. 1C** and **Table S1**), which are important for function (Feltner et al., 2016). No synonymous mutations in *lasR* were detected. Two mutations (Δ36-38 and L130P) were present in all three replicate cultures at Day 4 and thus were likely present in the initial inoculum. In replicate A, two additional mutations in *lasR* (E183G and A231V) were identified at Day 6; the LasR A231V substitution has been extensively characterized as a loss-of-function mutation through phenotyping and genetic complementation (Lujan, Moyano, Segura, Argarana, & Smania, 2007; Qi, Toll-Riera, Heilbron, Preston, & MacLean, 2016). The percentage of *lasR* mutants in the evolved population detected by sequencing at Day 4 (22.2 ± 6.0% s.d.) and Day 6 (48.5 ± 4.9% s.d.) (**Fig. 1C**) closely resembled the percentage of LasR^-^ strains predicted by the model (∼20% and ∼50%, respectively) (**Fig. 1A**). The increased frequency of cells with the allele encoding the L130P substitution (McCready, Paczkowski, Henke, & Bassler, 2019) between Day 4 and Day 6, with 13.1%, 24.6% and 41.4% increases in replicate cultures, suggests strong selection for this particular variant or the presence of an additional mutation(s) in this background. In support of the significant increases in LasR^-^ subpopulations, the evolved cultures themselves had lower levels of the LasR-regulated autoinducer 3OC12HSL; by Day 2, culture 3OC12HSL levels were ∼30% lower than a non-evolved wild-type culture, and showed a ∼90% reduction by Day 6 (**Fig. 1D**).

To further test the predictive power of our model for the rise of LasR^-^ lineages, we initiated cultures with different ratios of a constitutively tagged wild type (*att::lacZ*) against untagged wild-type or Δ*lasR* mutant competitors. A control assay demonstrated that the ratios of tagged and untagged wild type were unchanged over the course of growth, as shown previously (Clay et al., 2020; D. L. Mould et al., 2020). When the Δ*lasR* competitor was cultured with the tagged wild type, the percentage of Δ*lasR* mutant cells in the total population increased regardless of the initial percentage of Δ*lasR* (1% to 85%) at the time of inoculation (**Fig. 1E**), and the model successfully predicted the extent to which the Δ*lasR* would outcompete the wild type over this range (**Fig. 1E**-dotted line).

### Activity of CbrAB, the two-component system that regulates carbon utilization, is required for the rise of LasR^-^ strains

To test which genes or pathways were required to promote the selection of LasR^-^ cells, we applied reverse genetics to experimental evolution. In *P. aeruginosa,* the sensor kinases of two-component systems, encoded throughout the genome, respond to a variety of diverse internal and environmental cues, such as nutrient limitation or stresses, that may be relevant to differential fitness (Rodrigue, Quentin, Lazdunski, Méjean, & Foglino, 2000; B. X. Wang, Cady, Oyarce, Ribbeck, & Laub, 2021). Using a library of 63 sensor kinase deletion mutants (B. X. Wang et al., 2021), we screened each mutant for the rise of LasR^-^ phenotypes in triplicate in a 96-well plate format (**Fig. S3**). In the primary microtiter dish based screen, in which the investigators were blind to mutant strain identity, five gene knock-outs (*ΔcbrA,* Δ*gacS,* Δ*fleS,* ΔPA14_64580, and ΔPA14_10770) showed no detectable “sheen” colony phenotypes characteristic of LasR^-^ strains in any of the three replicates (**Fig. S3** & **Table S2**). In a secondary screen of these five mutants in five mL cultures, only the Δ*cbrA* mutant (**Fig. 2A**) did not evolve LasR^-^ phenotypes after serial passage; the other four mutants all had significant subpopulations with LasR-phenotypes by Day 6 (**Fig. S4A**). In addition, evolution experiments initiated with Δ*anr* or Δ*rhlR* mutants, which lack genes known to contribute to LasR^-^ strain phenotypes and fitness (Chen, Déziel, Groleau, Schaefer, & Greenberg, 2019; Clay et al., 2020), resembled wild type with at least 80% of colonies displaying LasR^-^ phenotypes by Day 8 (**Fig. S4B**).

**Figure 2.**
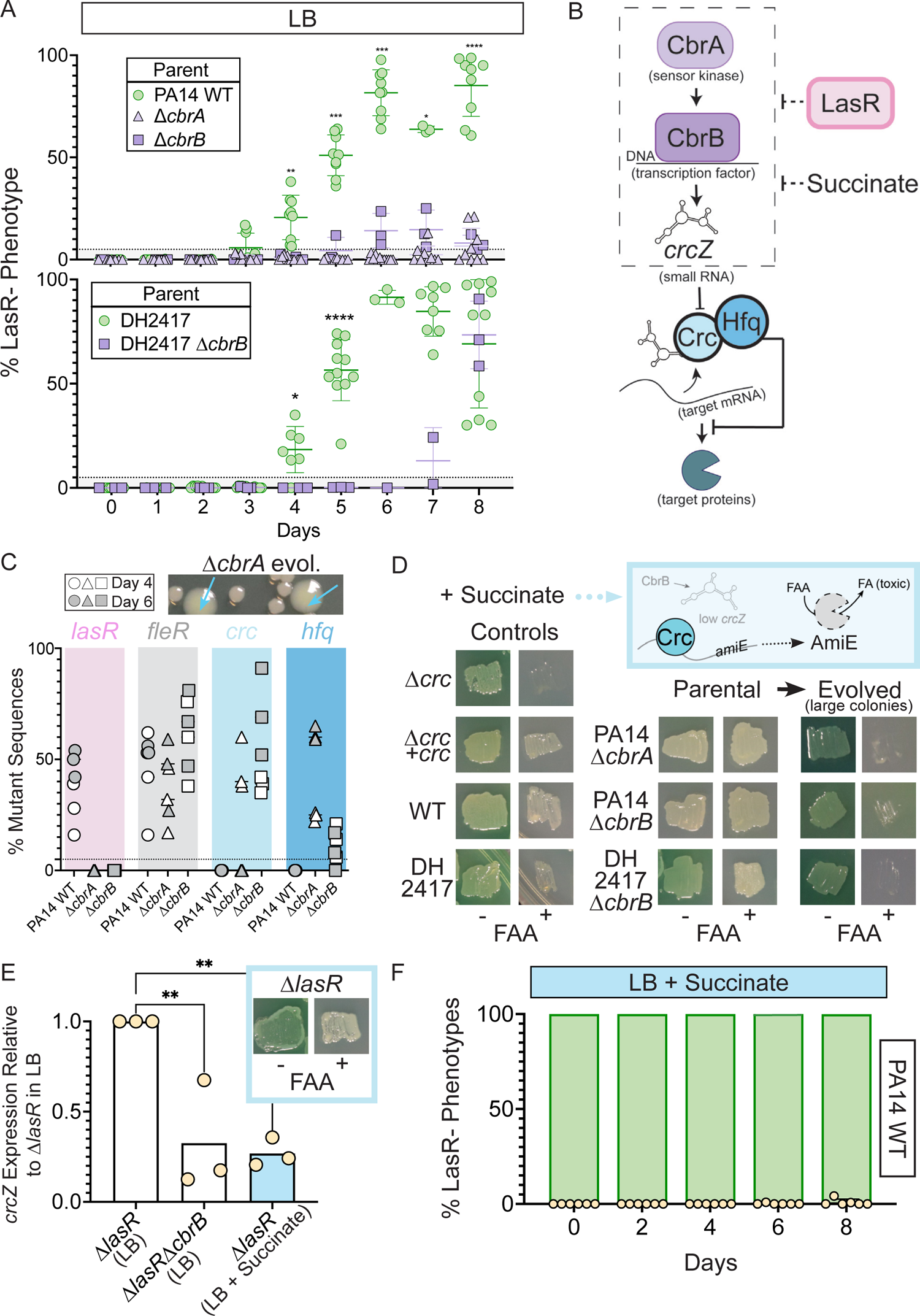
Activity of the carbon catabolite repression system is required for LasR-selection in LB. A. The percentage of colonies with LasR-phenotypes enumerated over the course of evolution for Δ*cbrA* or Δ*cbrB* mutants (purple triangle and square, respectively) in strains PA14 or a LasR+ cystic fibrosis isolate (DH2417) relative to “wildtype” comparators. PA14 WT strain data is the same as in Figure 1B (n 2: 3). Statistical significance determined between percent LasR-phenotypes in CbrA/B+ and *cbrA/B* mutant pools at each day via Two-Way AN0VA with Dunnet’s multiple hypothesis correction. *, p < 0.05; **, p < 0.005; ***, p < 0.0005; ****, p < 0.0001. B. The carbon catabolite repression system promotes the preferential consumption of succinate (and other preferred substrates) through the two-component system CbrAB. CbrA activates its response regulator CbrB which directly induces expression of the small RNA *crcZ*. *crcZ* sequesters Crc thereby allowing translation of the target gene to occur. 0ften the target gene enables the utilization of specific (i.e. less preferred) substrates. In a catabolite repressed state, such as when succinate is present, Crc binds to target mRNA with the RNA binding protein Hfq and blocks translation. CbrB protein levels are higher in strains lacking LasR function, but the mechanism linking these pathways is uncharacterized. C. Percent total mutant alleles in *lasR* (pink bar), *fleR* (grey bar), *crc* (light blue bar), and *hfq* (darker blue bar) in a representative experiment (Fig. 1B, diamond symbols) for PA14 wild type, Δ*cbrA*, Δ*cbrB* evolved populations sequenced on days four (white filled symbol) and six (grey filled symbol). Representative image of the larger colony morphologies observed in the evolved pools from CbrA/B-deficient strains (Δ*cbrA* shown) above. D. Crc represses *amiE* encoding an amidase that can turnover the luoroacetamide (FAA) protoxin to luoroacetate (FA) mediating cell death. ln the presence of succinate, cells with functional Crc survive in the presence of FAA. PA14 WT, PA14 Δ*cbrA*, PA14 Δ*cbrB*, and DH2417 WT strains were included as controls. The Δ*cbrA* and Δ*cbrB* parental strains used for the evolution experiments and representative colonies that emerged with a larger colony size in these backgrounds were patched (or struck out) onto succinate containing plates in the absence and presence of the FAA protoxin. E. *crcZ* expression of PA14 Δ*lasR* in LB (white bar) and LB with 40 mM succinate (blue bar) measured by qRT-PCR and plotted relative to expression of Δ*lasR* in LB (n = 4). lnset shows reperesntative image of Δ*lasR* grown on succinate containing plates in the absence and presence of FAA. F. Percentage of colonies with LasR-phenotypes observed in evolution experiment initiated with strains PA14 WT in LB supplemented with 40 mM succinate (n = 6).

CbrA, through its regulation of the response regulator CbrB, (D’Argenio, Wu, Hoffman, Kulasekara, Déziel, et al., 2007; E. Sonnleitner, Abdou, & Haas, 2009), controls *P. aeruginosa* preferential catabolism of certain carbon sources, such as succinate, over others (e.g. amino acids) through a process referred to as catabolite repression. In support of the finding that CbrA was essential for the evolution of LasR^-^ lineages, the Δ*cbrB* mutant also showed a striking and significant reduction in LasR^-^ phenotypes over the course of eight days (**Fig. 2A**). Additionally, evolution experiments in a LasR^+^ cystic fibrosis clinical isolate (DH2417) showed a similar rise in LasR^-^ phenotypes over the course of evolution, which was not observed in a Δ*cbrB* derivative (**Fig. 2A**). CbrAB-controlled catabolite repression is regulated by Crc, in complex with the RNA-binding protein Hfq, which together repress the translation of target mRNAs involved in the transport and catabolism of less preferred substrates (**Fig. 2B**) (Elisabeth Sonnleitner, Prindl, & Bläsi, 2017). Crc activity is down regulated by the small RNA *crcZ*, which sequesters Crc away from its mRNA targets. The CbrAB two component system transcriptionally regulates levels of *crcZ* (**Fig. 2B**) (E. Sonnleitner et al., 2009) in response to signals that have yet to be described.

Consistent with the absence of LasR^-^ phenotypes in evolved Δ*cbrA* or Δ*cbrB* cultures, Pool-Seq analysis found no mutations in *lasR* at either Day 4 or 6 (**Fig. 2C**, pink and **Table S1**) which was in striking contrast to the multiple LasR^-^ alleles observed in wild type cultures. The absence of *lasR* mutations in the Δ*cbrA* and Δ*cbrB* derivatives was not due to differences in mutation frequency or number of generations as other mutations in distinct pathways under selection (e.g. *fleR* in **Fig. 2C**) were present at comparable levels in all cultures (**Table S1** for data). In addition, strain PA14 wild type and the Δ*cbrA* mutant had similar growth patterns as assessed by daily optical density measurements (**Fig. S4C**). We also assessed a number of factors other than differential growth that could affect the rise of LasR^-^ lineages. A previous report (Heurlier et al., 2005) found that LasR^-^ strains in the PAO1 background undergo less severe alkaline-induced lysis in another complex medium (nutrient yeast broth) when grown aerobically, but we found no evidence of differential lysis in LB between wild-type and Δ*lasR* strains under our conditions (**Fig. S1A**). Furthermore, buffering the medium to pH 7 suppressed medium alkalinization (from pH of 6.8 to 8.5) and lysis (Crocker et al., 2019), but not the rise of LasR^-^ lineages; though, the kinetics of LasR^-^ lineage detection was delayed with buffering (**Fig. S4D** and (Sandoz et al., 2007)). Lastly, to assess potential differences in toxicity of the wild type and Δ*cbrB* mutant culture supernatants towards LasR^-^ cells through the production of secreted factors (Yan et al., 2018), we grew the Δ*lasR* mutant in spent filtrate from wild-type and Δ*cbrB* cultures; no significant differences in colony forming units were observed (**Fig. S4E**).

The activation of CbrAB increases growth on diverse metabolites by inducing *crcZ* which sequesters Crc away from the targets that it transcriptionally represses (**Fig. 2B** for pathway). In D’Argenio et al (D’Argenio, Wu, Hoffman, Kulasekara, Deziel, et al., 2007; D’Argenio, Wu, Hoffman, Kulasekara, Déziel, et al., 2007), higher CbrB levels were observed in LasR^-^ strains in a proteomics analysis, but no direct interactions between LasR and components of CbrA-CbrB-*crcZ*-Crc pathway have been described. Because CbrA, CbrB, and *crcZ* act to repress Crc, we hypothesized that if the loss of LasR function led to higher activity of the CbrA-CbrB-*crcZ* pathway and less Crc translational repression, we might also observe loss-of-function mutations in the genes encoding Crc or Hfq (**Fig. 2B**). Interestingly, the pooled genome sequence data from the Day 4 (open symbols) and Day 6 (grey symbols) populations evolved in the Δ*cbrA* and Δ*cbrB* backgrounds identified seven different mutations in *crc*, including three nonsense mutations, four missense mutations, and six indels, and these were among the most abundant mutations in the Δ*cbrB* mutant cultures; no *crc* mutations were identified in the PA14 wild type evolved populations (**Fig. 2C**). In Δ*cbrB*, *crc* mutant alleles showed the largest rise between Day 4 and Day 6 across all three replicate cultures (**Table S1**). In the Δ*cbrA* passaged cultures, we also identified a rise in *hfq* mutations within the coding and upstream intergenic regions (**Fig. 2C** and **Table S1** for sequence data) in addition to mutations in *crc*. The changes in relative abundances of alleles with mutations in *crc* and either the promoter or coding regions of *hfq* across the two days suggested that *hfq* mutations and *crc* mutations were in different backgrounds (**Table S1**).

To assess Crc-Hfq function in evolved strains, we leveraged Crc translational repression of the amidase AmiE, which cleaves the prototoxin fluoroacetamide (FAA) to the toxic fluoroacetate (FA) (**Fig. 2D** for pathway) (O’Toole, Gibbs, Hager, Phibbs, & Kolter, 2000). Succinate, which downregulates CbrAB activity, maintains repression of AmiE, thereby enabling wild type to grow in the presence of FAA. In the absence of functional Crc or its co-repressor Hfq, cells synthesize AmiE, and FAA conversion into FA inhibits growth. As expected, on medium with succinate, FAA inhibited growth of the Δ*crc* mutant, but did not affect growth of the complemented Δ*crc + crc* strain, the wild type and the Δ*cbrA* and *ΔcbrB* mutants. However, in passaged Δ*cbrA* and *ΔcbrB* cultures, spontaneous mutants in the population gave rise to larger colonies (**Fig. 2C**, top), and these isolates were FAA sensitive (**Fig. 2D**) supporting the model that in the Δ*cbrA* and Δ*cbrB* backgrounds, mutations that abolished Crc or Hfq activity arose.

Secondary mutants with FAA sensitivity also arose in the DH2417 Δ*cbrB* background upon passaging, indicating that this phenomenon was not unique to the PA14 background and another study also reported *crc* and *hfq* mutants in the absence of *cbrB* (Boyle et al., 2017). Given the apparent selection for decreased Crc function in Δ*cbrA* and *ΔcbrB*, and the requirement of *cbrA* or *cbrB* for LasR^-^ strain selection, we hypothesized that increased CbrAB activity may be a trait that increases the fitness of LasR^-^ strains.

To complement the genetics approach of evolution assays in *cbrAB* mutants, we monitored the rise of LasR^-^ lineages in LB medium supplemented with succinate, which inhibits CbrAB activity (E. Sonnleitner et al., 2009). Medium amendment with 40 mM (pH 7) succinate was sufficient to repress CbrB-regulated *crcZ* small RNA expression in Δ*lasR* to levels reminiscent of Δ*lasRΔcbrB* (**Fig. 2D**). The repression of *crcZ* in Δ*lasR* by succinate and Δ*lasR* growth on FAA + succinate, unlike Δ*crc* (**Fig. 2E**, inset), indicated that Δ*lasR* retains the control of Crc-Hfq mediated regulation. Succinate amendment suppressed the rise of LasR^-^ phenotypes in PA14 wild type (**Fig. 2F**).

### Elevated *cbrB* and *crcZ* expression and reduced Crc-dependent repression are sufficient to recapitulate the growth advantages of LasR^-^ strains

CbrAB activity induces the expression of *crcZ*, which sequesters Crc. We found that the Δ*lasR* mutant had ∼ two-fold higher *crcZ* levels compared to wild type, indicating higher activity of the CbrAB two component system in LasR^-^ strains (**Fig. 3A**). Previous work reported higher yields on phenylalanine for LasR^-^ relative to LasR^+^ strains concomitant with elevated CbrB protein levels in a proteomics analysis (D’Argenio, Wu, Hoffman, Kulasekara, Deziel, et al., 2007). Thus, we used phenylalanine along with other growth substrates to further dissect the activity of the CbrAB-*crcZ*-Crc pathway (**Fig. 2A**) in LasR^-^ strains. In planktonic cultures in medium with phenylalanine as a sole carbon source, the Δ*lasR* strain obtained significantly higher yields than the wild type and the enhanced growth phenotype was complementable by *lasR* (**Fig. 3B**). As previously reported, growth on phenylalanine depended on *cbrB*; the Δ*cbrB* and Δ*lasR*Δ*cbrB* mutants grew similarly poorly and their growth could be fully complemented by expressing *cbrB* (**Fig. 3B**). Deletion of *crc* in the Δ*lasRΔcbrB* strain also restored growth to levels comparable to the Δ*lasR* and Δ*lasRΔcbrB* + *cbrB* strains (**Fig. 3B**) indicating Crc repression of phenylalanine catabolism in the absence of CbrB. Overexpression of either *cbrB* or its target *crcZ*, which acts as a Crc-sequestering agent, was sufficient to improve yields on phenylalanine relative to the empty vector control (**Fig. 3C**). The CbrB- and Crc-controlled growth advantage on phenylalanine for LasR^-^ strains in planktonic cultures was also apparent in colony biofilms (**Fig. 3D**). One interesting difference between planktonic and biofilm assays was the differing requirement for CbrB for robust growth of wild type (**Fig. 3B** versus **3D**). On other substrates for which catabolism is under the control of Crc, e.g. mannitol and glucose, LasR^-^ strains showed complementable CbrB-dependent growth advantages over the wild type (**Fig. S5A**). While deletion of *crc* was able to restore enhanced growth to the Δ*lasR*Δ*cbrB* mutant, Δ*crc* did not grow as robustly as the Δ*lasR* mutant which is consistent with the detection of LasR^-^ lineages but not Crc^-^ lineages in passaged wild type cultures.

**Figure 3.**
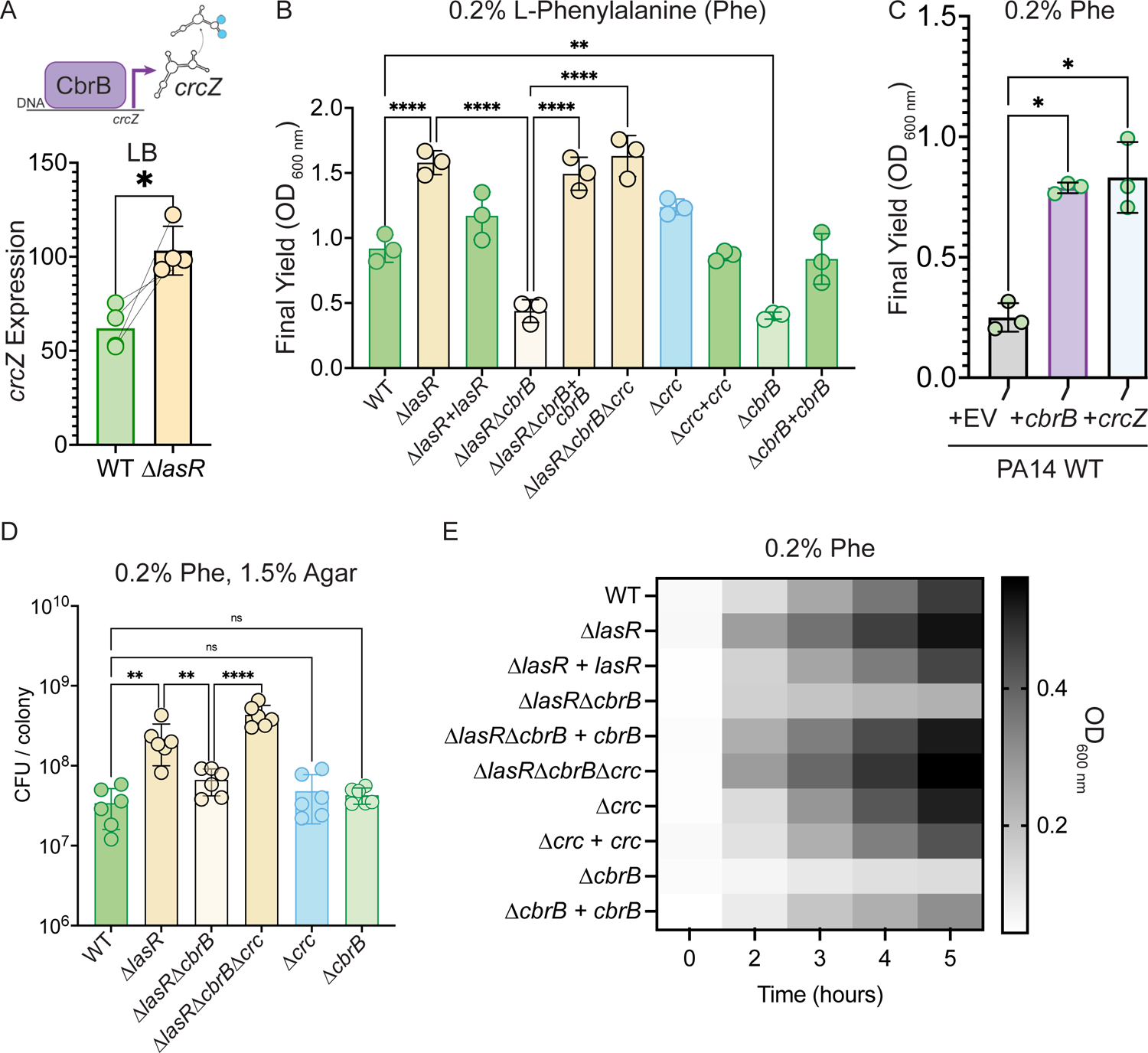
Increased CbrB activity of LasR-strains is necessary and sufficient to promote growth on non-repressive substrates like phenylalanine via Crc. A. CbrB promotes the transcription of *crcZ,* and *crcZ* is thus a direct readout of CbrB transcriptional activity. *crcZ* expression was measured by qRT-PCR relative to the average expression of the housekeeping genes *rpoD* and *rpsL* in cultures of PA14 WT and Δ*lasR* strains grown to OD_600 nm_ = 1 from four independent experiments. *, p < 0.05 as determined by Student’s t-test. B. Final yield on phenylalanine (Phe) as a sole carbon source shows enhanced growth for Δ*lasR, cbrB* dependence, and the requirement for *cbrB* is abolished by deletion of *crc*. Each point is the average of three replicates, repeated three independent days. Statistical significance determined by one-way AN0VA with Sidak’s multiple comparisons test. ns, not significant. *, p < 0.05. **, p < 0.005. ****, p < 0.0001. C. Final yield on Phe under (0.2%) arabinose-inducing conditions for the PA14 WT strain expressing an empty vector or *crcZ*, and *cbrB* overexpression constructs. Each point is the average of three replicates, performed on three separate days. Statistical signifcance determined by one-way AN0VA with multiple hypothesis correction. D. Colony counts (CFU) from resuspended colony biofilms grown on Phe as a sole carbon source for 24 h. Each point is a single replicate and the experiment was performed on six independent days. Statistical signifcance determined by one-way AN0VA with Sidak’s multiple comparisons test. E. Heatmap representation density during planktonic growth as measured by OD_600 nm_ on phenylalanine as a sole carbon source. n = 3. From same growth data used to generate Fig. 3B.

In addition to the higher yields relative to the wild type with phenylalanine as the sole carbon source, the Δ*lasR* strain also had a reduction in lag phase similar to what was observed in LB medium (**Fig. S1**). As shown in **Fig. 3E**, which displays kinetic data collected up to 5 hours from the experiments used to generate the data in **Fig. 2B**, Δ*lasR* or Δ*lasR*Δ*cbrB*Δ*crc* had markedly higher levels of growth by 2 hours after an LB-grown inoculum was transferred into fresh medium with phenylalanine as the sole carbon source, and higher Δ*lasR* densities persisted in exponential phase. The Δ*lasR*Δ*cbrB* mutant lacked this early growth enhancement and this defect was complementable. Thus, LasR^-^ strains from stationary phase cultures appear to be primed for growth on single carbon sources under CbrB-Crc control.

### LasR^-^ strains have CbrB-dependent growth advantages on metabolites enriched in progressive cystic fibrosis lung infections

Loss-of-function mutations in *lasR* are commonly detected in samples from chronic *P. aeruginosa* lung infections in pwCF, and these mutants have been correlated with a more rapid rate in lung function decline (Hoffman et al., 2009). To determine the metabolite milieu in the CF lung, we performed a metabolomics analysis of bronchioalveolar lavage samples collected from ten pwCF and ten non-CF individuals (**Table S3**). The pwCF were infected with diverse pathogens and had varying lung function, which was measured as forced expiratory volume in one second and presented as the percent expected at one’s age (%FEV_1_). Over 300 compounds were measured, and no uniquely microbial metabolites were noted. Many compounds were higher in the CF population, but some were unchanged (e.g. glucose) and others were higher in non-CF samples (e.g. adenosine and glutathione as previously published (Esther et al., 2008; Fitzpatrick, Park, Brown, & Jones, 2014) (**Table S4**).

In a principal component analysis (PCA), samples from non-CF individuals clustered together while those from pwCF were more spread. Samples from pwCF with high lung function (112 or 113 %FEV_1_) grouped among the non-CF samples (**Fig. 4A**). The metabolites that contributed strongly to the first principal component, PC1, showed a significant inverse correlation with %FEV_1_ including phenylalanine, arginine, lactate and citrate. As for phenylalanine (**Fig. 3B** & **D**), the Δ*lasR* strain had growth advantages on arginine, lactate, and citrate that were controlled by CbrB and Crc (**Fig. S5B,C**).

**Figure 4.**
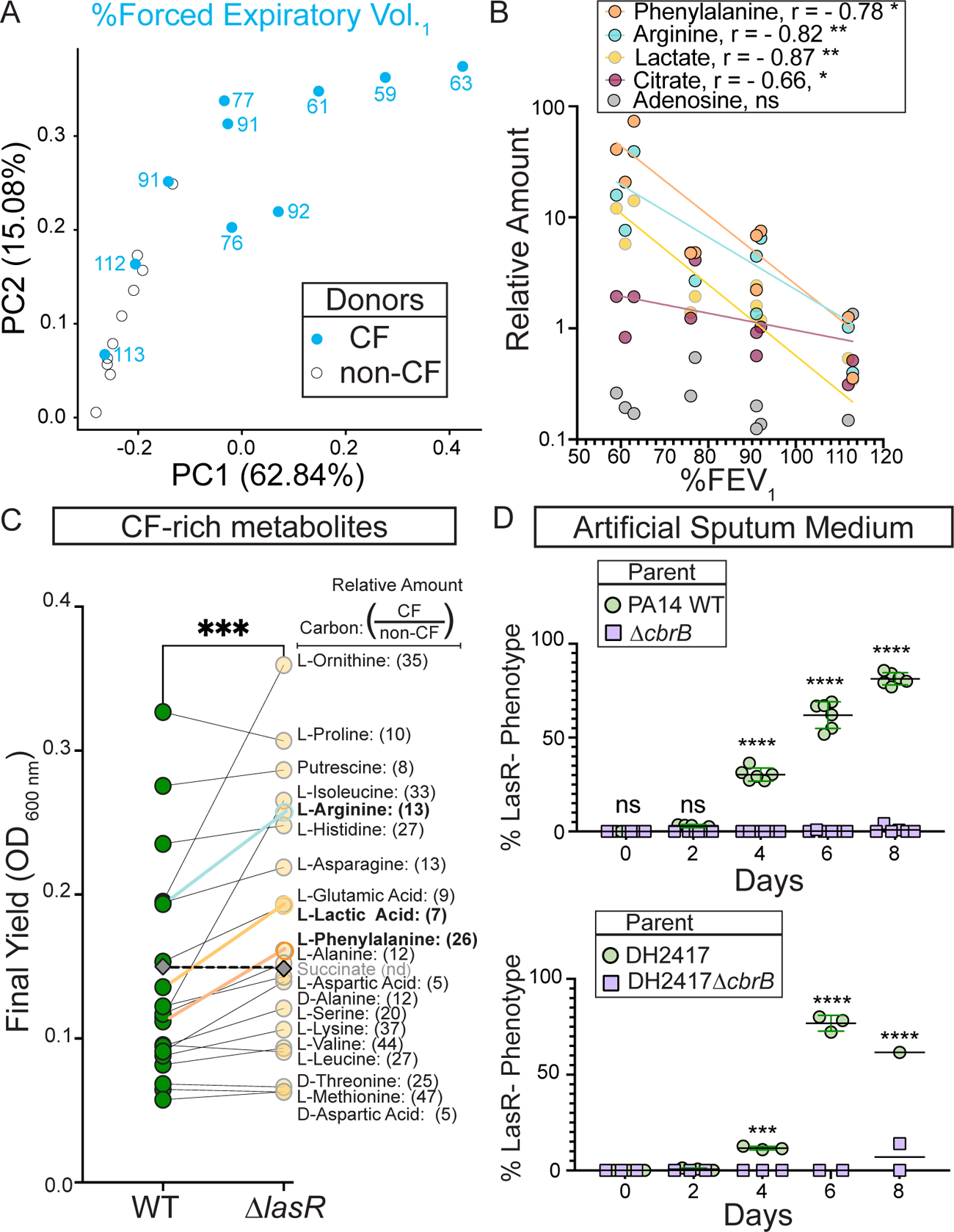
CbrB-dependent growth advantages may contribute to *lasR* mutant selection in distinct nutrient profiles of progressive cystic fibrosis airways. A. The first two dimensions (PC1 and PC2) of a principal component analysis of log normalized metabolite counts from bronchoalveolar lavage (BAL) samples collected from cystic fibrosis (CF, blue) and non-cystic fibrosis (non-CF, grey) donors explain 62.84% and 15.08% of the variation in the data, respectively. PC1 separates the metabolite data by relative lung function as measured by percent forced expiratory volume in 1 sec (%FEV_1_) for samples from people with CF. The %FEV_1_ is overlayed for CF-donor samples with text. Samples from non-CF donors group more closely with CF-donor samples that have high lung function. B. Spearman correlation analysis of the relative phenylalanine (orange), arginine (aqua), lactate (yellow), and adenosine (grey) metabolite counts in the BAL samples relative to %FEV_1_. C. Final yield measured after 24 h for strains PA14 WT and Δ*lasR* on a subset of carbon sources in Bl0L0G growth assays for which the metabolite was found to be in higher abundance in CF-donor relative to non-CF donor BAL samples. Bold font indicates carbon sources analyzed in Fig. 3 and Supp. Fig. 5. Number in parenthesis refers to the ratio of the average counts for each metabolite in CF relative to non-CF samples. D. 0bserved percentage of colonies with LasR-phenotypes over the course of evolution from strains (top) PA14 WT or (bottom) CF isolate (both green cirlces) with Δ*cbrB* (purple squares) derivatives in artificial sputum medium (ASM), which was designed to recapitulate the CF lung nutritional profile. ***, p = 0.0008; ****, p < 0.0001 as determined by ordinary two-way AN0VA with Sidak’s multiple comparisons test.

We identified the twenty carbon sources that were most enriched in CF samples including those that correlated inversely with lung function, then used a BIOLOG phenotype array to assess whether the trend of greater yield for the Δ*lasR* strain persisted across this set. We confirmed that the Δ*lasR* strain reached significantly higher yields on the metabolites including phenylalanine, arginine, and lactate and that overall Δ*lasR* showed better growth than the wild type (**Fig. 4C**).

To further test the hypothesis that LasR^-^ strains evolved due to enhanced growth in the nutrient environment of the CF lung, we performed evolution experiments using both strain PA14 and a LasR^+^ CF clinical isolate in a medium designed to more closely recapitulate the nutritional profile of the cystic fibrosis airway. Upon absolute quantitation, we observed good concordance between the relative abundances of amino acids found in BAL fluid and reported for sputum (Palmer, Mashburn, Singh, & Whiteley, 2005) which served as a basis for an artificial sputum medium, ASM (**Fig. S6**) (Clay et al., 2020) which was based on a previously reported synthetic CF medium (SCFM2) (Palmer et al., 2005). LasR^-^ strains evolved in both strain backgrounds (**Fig. 4D**) with kinetics similar to what was observed in LB medium (**Fig. 1B**). Parallel evolution experiments in ASM initiated with Δ*cbrB* derivatives did not exhibit a rise in LasR^-^ phenotypes in either strain background to suggest that CbrAB activity was again a contributor to the fitness of *lasR* loss-of-function mutants.

## Discussion

Through mathematical modeling, experimental evolution and competition assays, we found that the rise of problematic *P. aeruginosa* LasR^-^ variants frequently observed in disease could be explained by increases in yield and decreases in lag during growth on carbon sources abundant in the lung environment (**Fig. 5**). In fact, the steady state growth rate for Δ*lasR* was slightly less than that for the wild type, which is consistent with the model that there are frequently tradeoffs between a shorter lag phase and overall growth rate (Basan et al., 2020). Interestingly, CF-adapted *P. aeruginosa* isolates have been found to have slower *in vitro* growth rates than other strains (Yang et al., 2008). Other factors will impact the relative fitness of LasR^+^ and LasR^-^ cells across different growth phases (**Fig. 5**) including oxygen availability and pH buffering capacity, which may lead to differential lysis (Heurlier et al., 2005), or the need for (or exploitation of) proteases to gain access to growth substrates (Barbieri, Delden, Pesci, Pearson, & Iglewski, 1998; Sandoz et al., 2007).

**Figure 5.**
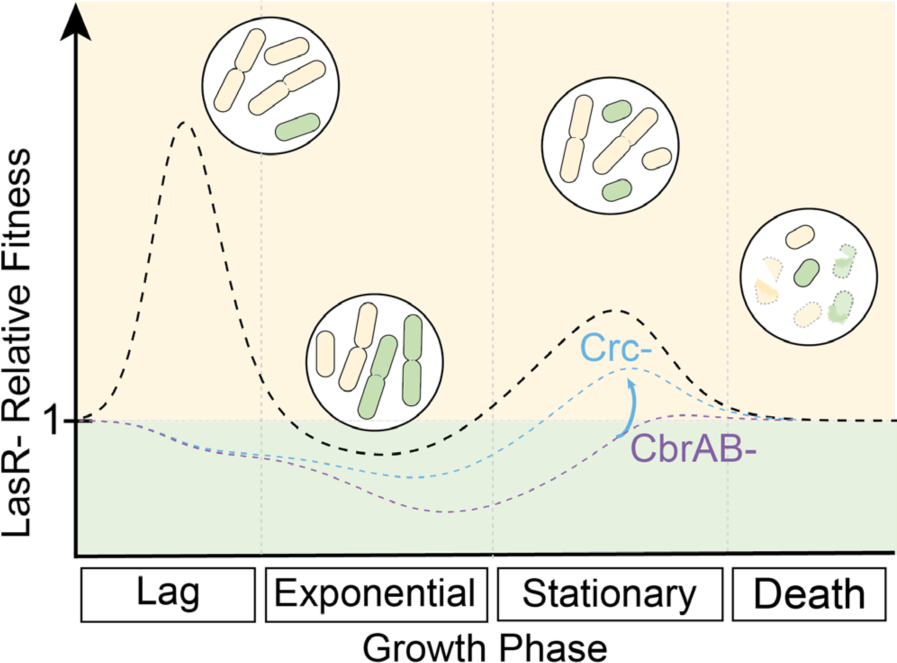
CbrAB activity contributes to the positive selection of LasR-strains in complex media. LasR-strain fitness relative to wild type is shown across growth phases, including lag, exponential growth, stationary, and death phases. Relative fitness of LasR-strain (dotted black line) is calculated from the experimentally determined monoculture growth data of strains PA14 wild type (WT) and Δ*lasR* over time. Values above one indicate a LasR-strain fitness advantage over the WT strain during that growth phase. Circled insets show representative cartoons of LasR-(beige) and LasR+ (green) cells at each growth phase to indicate dividing or lysing cells (burst cells) across growth stages. The heights of the peaks or valleys of the relative fitness lines can be altered by several modulating factors including those that contribute to the positive and negative selection of LasR-strains. Other modulating factors reported or suggested in the literature include inter- and intra-species competition, extracellular protease, immunoclearance, and oxygenation which are likely condition dependent. In the absence of CbrA or CbrB (CbrAB-, dotted purple line) or in the presence of succinate (one CbrAB repressive substrate), the relative fitness of LasR-strains is lower resulting in a reduction in the observed selection. This could be partially relieved in the CbrAB-background through disruption of Crc or Hfq function (blue dotted line), restoring activity through the pathway.

Data presented support the model that that increased growth of LasR^-^ cells on many amino acids, sugars and lactate, is due to higher CbrAB-controlled *crcZ* levels which downregulates metabolism under Crc control, and these findings nicely parallel studies by D’Argenio et al. (D’Argenio, Wu, Hoffman, Kulasekara, Déziel, et al., 2007) that found higher levels of CbrB in LasR^-^ isolates. In Δ*cbrA* and Δ*cbrB* mutants, *lasR* loss-of-function mutations did not arise, but mutations in *crc* and upstream of *hfq* were observed. As *crc* mutations phenocopy many of the growth advantages of the *lasR* mutants (**Fig. 2C**, **Fig. 3**, **Fig. S5**), the importance of derepressed catabolism for fitness is underscored. Though deletion of *cbrA* or *cbrB* can have pleiotropic effects (Yeung, Bains, & Hancock, 2011), we did not observe differences in density, quorum sensing regulation and production of quorum sensing controlled factors such as proteases, lysis in stationary phase, or overall mutation accumulation between wild type, Δ*cbrA*, and Δ*cbrB* that could explain differences in the rise of LasR^-^ subpopulations. Furthermore, environmental modification of CbrB activity by the addition of succinate to LB (E. Sonnleitner et al., 2009) also suppressed the emergence of LasR^-^ strains in the wild type. Because CbrAB activity can still be suppressed by succinate in LasR^-^ cells (**Fig. 2E**), LasR^-^ variants were not strictly “de-repressed”, and this is consistent with the fact that Δ*lasR* and Δ*crc* growth patterns were not identical. Unlike *lasR* mutations, *crc* mutations are not commonly observed in clinical isolates (Winstanley, O’Brien, & Brockhurst, 2016) and thus are likely also under negative selection despite some growth advantages (Lorenz et al., 2019). Several CF isolates show reduced succinate assimilation to suggest the uptake of less preferred substrates over the course of adaptation (Jørgensen et al., 2015; La Rosa, Johansen, & Molin, 2018).

Analysis of BAL fluid revealed higher levels of substrates such as lactate and amino acids, which require CbrB for consumption in samples from pwCF, and these findings are consistent with other more targeted analyses of CF airway samples (Bensel et al., 2011; Twomey et al., 2013). Consistent with our finding that higher levels of certain metabolites correlated with worse CF lung disease, other studies including that of Esther et al. (Esther et al., 2016) found a correlation between total metabolites and neutrophil counts suggesting host cell lysis, along with lysis of microbial cells, may be a major contributor to a shift in the metabolome. CF-lung derived *P. aeruginosa* isolates can have amino acid auxotrophies and enhanced amino acid uptake (La Rosa, Johansen, & Molin, 2019) which supports ready access to amino acids *in vivo*.

Our model predicts LasR^-^ strains benefit from growth advantages that might be present when new nutrients become available (analogous to lag phase) and in dense populations when improved yields for the Δ*lasR* mutant emerges; due to a slower steady-state growth rate, we predict that LasR^-^ strains would not emerge under steady-state growth conditions such as in a chemostat. Indeed, the advantages of decreased lag phase in cultures, even at the expense of steady-state growth rates, has been proposed to be a universal adaptation in dynamic environments (Basan et al., 2020). Thus, the frequent emergence of LasR^-^ lineages in the CF lung and other disease settings suggests that *P. aeruginosa* often undergoes growth transitions *in vivo*, possibly due to fluctuating local conditions, spatial heterogeneity, or the result of complex competition between bacterial and host cell types. In addition, the loss of LasR function enables other inherent advantages that contribute to competitive fitness including resistance to lysis under conditions of high aeration, enhanced microoxic fitness, enhanced RhlR activity (Chen et al., 2019; Clay et al., 2020; D’Argenio, Wu, Hoffman, Kulasekara, Deziel, et al., 2007; Heurlier et al., 2005); the connection between these phenotypes and the CbrAB-*crcZ*-Crc pathway is not yet clear.

The increased growth in post-exponential phase cultures for LasR^-^ strains bears similarities to mutations that arise in other microbes. For example, the selection for *rpoS* mutants in stationary phase cultures of *E. coli* (Finkel & Kolter, 1999; Zambrano, Siegele, Almirón, Tormo, & Kolter, 1993; Zinser & Kolter, 2000) is also dependent on nutrient accessibility (Farrell & Finkel, 2003) with enhanced amino acid catabolism as a major contributor to *E. coli* lineages with growth advantages in stationary phase (GASP) (Zinser & Kolter, 1999). While the rise of *rpoS* mutants in laboratory settings required pH-driven lysis (Farrell & Finkel, 2003), LasR^-^ strains still evolved in buffered medium suggesting distinct mechanisms for the metabolic advantages of *lasR* mutants. It is worth noting that none of the common GASP mutations (*rpoS, lrp*, or *ybeJ-gltJKL*) were identified in our *in vitro* evolution studies (**Table S1**). We considered that the enhanced growth of LasR^-^ strains in post-exponential growth phases may be due to differences in ppGpp signaling, given growth arrest as part of the stringent response modifies the expression of QS-regulated genes (van Delden, Comte, & Bally, 2001). However, no mutations in *relA* or *spoT*, the two ppGpp synthases, were observed. The mechanism of increased CbrB activity in Δ*lasR* remains an unresolved question that is relevant to *P. aeruginosa* biology and may aid in the identification of the signals that activate the CbrA sensor kinase which influences clinically-relevant phenotypes including virulence and antibiotic resistance (Yeung et al., 2011). Our working model is that the upregulation of CbrB transcription of *crcZ* increases levels of transporters and catabolic enzymes due to the release from Crc repression, and this enhanced substrate uptake alters intracellular metabolite pools driving metabolism in accordance with Le Chatelier’s principle (Monod, 1949). Thus, quorum sensing mutants can maintain higher growth rates at lower substrate concentrations than for quorum-sensing intact cells.

The repeated observation that *lasR* loss-of-function mutations readily arise in diverse settings provokes the question of how quorum sensing is maintained. Several elegant mechanisms that address this point have been described. First, the wiring of the LasR regulon is such that while there are growth advantages on many substrates present in the lung, there are growth disadvantages on other important nutrient sources (e.g. adenosine and proteins and peptides (Heurlier et al., 2005)). Social cheating can promote the rise of *lasR* loss-of-function mutants in protease-requiring environments (Diggle et al., 2007; Hassett et al., 1999). Second, there are quorum sensing controlled “policing” mechanisms through which LasR^+^ strains restrict the growth of LasR^-^ types through the release of products toxic to quorum-sensing mutants (Castañeda-Tamez et al., 2018; Rodolfo García-Contreras et al., 2020; M. Wang et al., 2015). Lastly, there are other tradeoffs such as sensitivity to oxidative stress that may limit LasR^-^ lineage success (Hassett et al., 1999). Quorum sensing exerts metabolic control in other diverse microbes beyond *P. aeruginosa*. Thus, these data provide insight into generalizable explanations for the benefits of metabolic control in dense populations and indicate drivers for frequent loss-of-function mutations in quorum-sensing genes such *agr* mutations in *Staphylococcus aureus* and *hapR* mutations in *Vibrio cholerae* (Dallas L. Mould & Hogan, 2021).

Together, these data highlight the power of coupling *in vitro* evolution studies with forward and reverse genetic analyses. Other benefits to this approach include the ability to dissect subtle differences between pathway components. For example, multiple mutations in *crc* repeatedly rose in Δ*cbrA-*, but not in Δ*cbrB*-derived populations, and multiple mutations in *hfq* rose in Δ*cbrB*-, and not in Δ*cbrA*-derived populations. While CbrA and B work together as do Crc and Hfq, these observations may provide a foothold into key distinctions that could yield mechanistic insights. In the future, the ability for deep sequencing of infection populations and analysis of evolutionary trajectories may aid diagnoses and treatment decisions in beneficial ways.

## Methods

### Strain Construction and Maintenance

In-frame deletions and complementation constructs were made using a *Saccharomyces cerevisiae* recombination technique described previously (Shanks, Caiazza, Hinsa, Toutain, & O’Toole, 2006). The *cbrB* and *crcZ* expression vectors were constructed by HiFi Gibson assembly according to manufacturer’s protocol. All plasmids were sequenced at the Molecular Biology Core at the Geisel School of Medicine at Dartmouth. In frame-deletion and complementation constructs were introduced into *P. aeruginosa* by conjugation via S17/lambda pir *E. coli*. Merodiploids were selected by drug resistance and double recombinants were obtained using sucrose counter-selection and genotype screening by PCR. Expression vectors were introduced into *P. aeruginosa* by electroporation and drug selection. All strains used in this study are listed in **Table S6**. Bacteria were maintained on lysogeny broth (LB) with 1.5% agar. Yeast strains for cloning were maintained on YPD (yeast extract-peptone-dextrose) with 2% agar. Artificial sputum medium (ASM) was made as described previously (Clay et al., 2020).

### Mathematical Model

Growth parameters were determined from 5 mL grown LB cultures inoculated as described in the experimental evolution protocol. Using a plate reader, the density (OD_600 nm_) was measured by taking a 100 µL aliquot at the designated time intervals with 1:10 dilutions for values greater than one. Lag and growth rate were measured in separate experiments from those used to monitor lysis. See extended note on mathematical model and Matlab script for additional details.

### Experimental Evolution

Experimental evolution was modeled after work by Heurlier et al. (Heurlier et al., 2005). A single colony of each strain was used to inoculate a 5 mL LB culture in 13 mm borosilicate tubes. The tubes inoculated with a single colony were grown for 24 h at 37°C on a roller drum. The 24 h grown culture was adjusted to OD_600 nm_ = 1 based on OD_600 nm_ reading of a 1 to 10 dilution in LB of the 24 h culture in a 1 cm cuvette using a Spectronic GENESYS 6 spectrophotometer. Separate 250 µL aliquots of the OD_600 nm_ normalized cells was sub-cultured into three tubes containing 5 mL fresh media to initiate the evolution experiment (i.e. time 0) with three distinct replicate cultures per experiment. At time of passage every two days, 25 µL of culture was transferred into 5 mL fresh media. Every day (or as indicated) cultures were diluted and bead spread onto LB agar plates for phenotype distinction. The LB agar plates were incubated for ∼24 h at 37°C and then left at room temperature for phenotype development. The sheen LasR^-^ colony morphologies were counted, and the percentage calculated based on total CFUs. All experimental evolutions in LB were repeated on at least three independent days with three replicates of each strain per experiment unless otherwise stated. The ASM and succinate amended medium evolutions were completed on two separate days. In the case of Δ*rhlR* and *Δanr*, the three replicates were inoculated from three independent overnights. Data visualization and statistical analysis was performed in GraphPad Prism 9 (version 9.2.0).

### gDNA extraction, Sequencing, SNP calling of Pool-Seq data

Between 100 and 150 random colonies were scraped and pooled from the LB agar plates that were counted and used to measure the percent of colonies with LasR^-^ phenotypes at days four and six from a representative WT-, Δ*crbA-*, and Δ*cbrB-*initiated evolution experiment. For plates containing a total of 100 ^-^ 150 colonies, all colonies on the plate were collected for a single pooled genomic DNA extraction. If more than 150 colonies were on a plate, the plate was divided equally, and all colonies in an arbitrary section were collected to ensure genomic DNA was extracted from a similar number of colonies for each sample. Scraped up cells were pelleted briefly in a 1.5 mL Eppendorf tube via a short spin, resuspended in 1 mL PBS, vortexted briefly, and gDNA was subsequently extracted from a 50 µL aliquot of cell resuspension via the Master Pure Yeast DNA purification kit according to manufacturer’s protocol with RNAase treatment. A 2.5 µg aliquot was submitted for Illumina sequencing (1Gbp) at the Microbial Genome Sequencing (MiGs) Center on the NextSeq 2000 platform. The resulting forward and reverse reads were trimmed. Both forward and reverse read files were aligned and compared to the complete and annotated UCBPP-PA14 genome available on NCBI (accession GCF_000014625.1) using the variant caller BreSeq (Deatherage & Barrick, 2014) (version 0.35.4) with a 5% cutoff. Specifically, the following command was used: breseq -r [reference file] [sample name]_.fastq.gz [sample name]_fastq.gz -o [output file name]. This provided an output file that specified variations from the reference genome and listed their respective fractions of the total reads. These fractions were treated as estimations of genotype proportions in the population. Variants at fixation (100%) across all 18 samples (three strains, two days) were excluded from follow-up analysis as potential differences in strain background that differed from the reference genome at the start of the experiment. All sequencing data is available on the Sequence Read Archive with accession number PRJNA786588 upon publication.

### Milk Proteolysis

Brain Heart Infusion Agar was supplemented with powdered milk dissolved in water to a final concentration of 5%. The evolved isolates selected on basis of “sheen” colony morphology were grown in a 96-well plate with 200 µL LB per well for 16 h. Milk plates were inoculated with ∼5 µL of culture using a sterilized metal multiprong inoculation device (Dan-Kar) and incubated at 37°C for 16 h. PA14 WT and Δ*lasR* strains were included as controls. Colonies which showed a halo of clearing larger than the Δ*lasR* control strain were considered protease positive.

### Acyl Homoserine Lactone Autoinducer Bioreporter Assays

Protocol as described in (D. L. Mould et al., 2020). Briefly, 100 µL of OD_600 nm_ normalized LB overnight cultures (OD_600 nm_ = 0.01) of the AHL-synthesis deficient reporter strains DH161 (3OC12HSL-specific) or DH162 (3OC12HSL or C4HSL responsive) with AHL-responsive promoters to *lacZ* (Whiteley & Greenberg, 2001; Whiteley, Lee, & Greenberg, 1999) were bead spread on LB plates containing 150 µg/mL 5-bromo-4-chloro-3-indolyl-β-D-galactopyranoside (XGAL, dissolved in DMSO). Inoculated plates were allowed to dry 10 min in a sterile hood. Once dry, 5 µL of either the test strains or control cultures (PA14 wild type and Δ*lasR* strains) were spotted onto the inoculated reporter lawns. After the spots dried, plates were incubated at 37°C for 16 h then stored at 4°C to allow for further color development, if necessary, based on wild-type colony activity. The blue halo that formed around the colony was interpreted as AHL activity. The levels of AHL produced are approximated by the size of the blue halo formed around the colony.

### Competition Assays

Competition assays were performed by competing strains against an *att::lacZ* strain as previously reported (Clay et al., 2020). Overnight cultures of *att::lacZ* competitor and test strains were normalized to OD_600 nm_ = 1 and mixed in the designated ratios with either a wild type control or Δ*lasR* strain. Aliquots of 10^-6^ dilutions of the initial mixed inoculums were immediately plated on LB plates containing 150 µg/mL XGAL by spreading an aliquot of 25 - 50 µL with sterilized glass beads. Roughly 100 - 200 colonies were counted to determine the initial ratios of PA14 *att:lacZ* to Δ*lasR* or the WT control strains by blue:white colony phenotype, respectively. To begin the competition experiment, a 250 µL aliquot of each undiluted mixed inoculum was sub-cultured into 5 mL fresh LB medium and incubated on a roller drum at 37°C for 6 h. After 6 h, the cultures were collected, diluted by 10^-6^ in fresh liquid LB, and plated as previously stated for blue:white colony screening. The LB plates containing XGAL were incubated overnight at 37°C prior to counting. Competitions were repeated on three separate days.

### Kinase Mutant Evolution Screen

Using an ethanol/flame sterilized metal multiprong inoculation device (Dan-Kar), the kinase mutant library (B. X. Wang et al., 2021) was inoculated into a 96-well plate with 200 µL LB per well for 24 h shaking at 37°C. The 24 h grown cultures were used to inoculate two 96-well plates with each kinase mutant (including PA14 WT control) in triplicate. These cultures were grown for 48 h upon which 2 µL was transferred to new 96 well plates with fresh 200 µL LB liquid per well. Each day, the wells containing the wild-type replicates were diluted by 10^-6^ in fresh LB and 25 µL was bead spread onto LB for phenotypic distinction based on sheen colony morphology. At day 14, when all wildtype replicates contained at least 50% LasR^-^ phenotypes, all wells were diluted and plated as stated previously for determination of sheen colony morphology. A secondary screen in 5 mL LB (as described above) was initiated for those mutant strains which did not show any LasR^-^ phenotypes across all three replicates. The Circos plot summarizing the screen data was generated using BioCircos (Cui et al., 2016) in R (version 4.0.2) and re-colored in Adobe Illustrator.

### Filtrate Toxicity

Based on a protocol used previously (Abisado et al., 2021), strains were grown 16 h in LB (5 mL) on a roller drum at 37°C, centrifuged at 13K RPM for 10 min in 2 mL aliquots and the resulting supernatant was filter sterilized through a 0.22 µm pore filter. Per 5 mL filtrate, 250 µL of fresh LB was added. A 16 h grown LB culture (5 mL) of PA14 Δ*lasR* was normalized to an OD_600 nm_ = 1, and 250 µL was used to inoculate 5 mL of the filtrate-LB mixture. The Δ*lasR* cultures were grown for 24 h at 37°C on the roller drum upon which colony counts were determined by bead spreading an appropriate dilution on LB plates. Data visualization and statistical analysis were performed in GraphPad Prism 9 (version 9.2.0).

### Fluoroacetamide Sensitivity Assay

Strains were inoculated (either by patching from plates or by spotting 5 µL of 16 h LB grown culture) onto plates containing 1.5% agar with M63 salts,10 mM lactamide, and 40 mM succinate with or without 2.5 mg/mL filter sterilized fluoroacetamide (FAA) dissolved in water based on protocol by Collier et al (Collier, Spence, Cox, & Phibbs, 2001). Relative growth was compared in the presence and absence of FAA. PA14 wild type and Δ*crc* were included as controls in every experiment wherein wild type displays robust growth on FAA in the presence of succinate and the Δ*crc* strain, little to none.

### Quantitative RT-PCR

The indicated strains were grown from single colonies in 5 mL LB cultures on a roller drum for 16 h, normalized to an OD_600 nm_ of 1 and 250 µL of normalized culture was inoculated into 5 mL fresh LB for a starting inoculum around OD_600 nm_ = 0.05. The cultures were then grown at 37°C on a roller drum until OD_600 nm_ = 1 at which point a 1 mL aliquot of culture was pelleted by centrifugation for 10 min at 13K RPM. Supernatant was removed and the cell pellets were flash frozen in an ethanol dry ice bath. This was repeated on three separate days with one WT and one Δ*lasR* culture pair (n = 4) collected on each day or one Δ*lasR* and one Δ*lasRΔcbrB* culture pair (n = 3) each day. Pellets were stored at −80°C until all sets of pellets were collected. RNA was extracted using the QIAGEN RNAeasy kit according to the manufacturer’s protocol, and 7 μg RNA was twice DNAse treated with the Turbo DNA-free kit (Invitrogen). DNA contamination was checked by semi-quantitative PCR with gDNA standard for 35 cycles with *rpoD* qRT primers; if DNA contamination was greater than 0.004 ng / μL, the sample was DNAse treated again. cDNA was synthesized from 400 ng of DNase-treated RNA using the RevertAid H Minus first-strand cDNA synthesis kit (Thermo Scientific), according to the manufacturer’s instructions for random hexamer primer (IDT) and a GC-rich template alongside an NRT control. Quantitative RT-PCR was performed on a CFX96 real-time system (Bio-Rad), using SsoFast Evergreen supermix (Bio-Rad) according to the following program: 95°C for 30 s and 40 cycles of 95°C for 5 s and 60°C for 5 s followed by a melt curve with 65°C for 3 s up to 95°C in increments of 0.5°C. Transcripts were normalized to the average *rpoD* and *rspL* expression unless stated otherwise. *rspL* and *crcZ* primers as designed in (Xia et al., 2020). *rpoD* primers as designed in (Harty et al., 2019). Data visualization and statistical analysis performed in GraphPad Prism 9 (version 9.2.0).

### Mono-carbon Growth

Single carbon sources were supplemented into M63 base (Neidhardt, Bloch, & Smith, 1974) and filter sterilized. A 16 h overnight LB culture grown at 37°C on a roller drum was normalized to an OD_600 nm_ = 1 in 2 mL LB. For liquid growth curves, a 250 µL aliquot of the density adjusted culture was spiked into 5 mL fresh M63 medium with designated carbon source in triplicate and growth was monitored using a Spec20 hourly in 13 mm borosilicate tubes. Every point on the liquid growth heat maps is the average of 3 replicates per day, repeated 3 days total. For colony biofilm growth, 5 µL of OD_600 nm_ normalized culture was inoculated onto 1.5% agar plate of M63 medium containing the designated carbon source in singlicate and grown for 16 h at 37°C. Colonies were cored using the back of a P1000 tip and disrupted by 5 min on Genie Disrupter in 1 mL LB. Disrupted colony biofilms were serially diluted. 5 µL of the serial dilutions were plated and a 50 µL aliquot of diluted colony resuspension (10^-6^ or 10^-7^-fold, depending on condition/strain) was bead spread and counted for colony forming units. Colony biofilm growth was assessed on > 5 independent days. Data visualization and statistical analysis performed in GraphPad Prism 9 (version 9.2.0).

### Metabolomics of Bronchioloalveolar Lavage Fluid and Artificial Sputum Medium

Human samples from people with and without cystic fibrosis were obtained with informed consent following institutional review board-approved protocols at Geisel School of Medicine at Dartmouth. The investigators were blinded to the conditions of the experiments during data collection and analysis. To obtain relative metabolite counts, bronchioloalveolar lavage (BAL) fluid samples were briefly centrifuged to exclude large debri then the supernatant was flash frozen in liquid nitrogen. Samples were processed by Metabolon via LC/MS for relative metabolite amounts. Raw values from Metabolon were normalized to protein concentrations by the BioRad Bradford protein concentration or raw area counts per day sample run and then the values were rescaled to set the median to 1. Missing values were imputed with the minimum rescaled value for that biochemical. Quantitative amino acid concentrations were determined for aliquots of the same BAL samples (lyophilized) using the Biocrates AbsoluteIDQ p180 kit at the Duke Proteomics Core Facility. The lyophilized samples of BAL were homogenized in water and 50/50 water/methanol respectively to extract metabolites. 25 µL of the BAL extract were utilized for preparation of the samples on a Biocrates AbsoluteIDQ p180 plate. A Waters Xevo-TQ-S mass spectrometer was utilized to acquire targeted metabolite quantification on all samples and quality control specimens. Raw data (in µM) was exported independently for the FIA-MS/MS and UHPLC-MS/MS acquisition approaches used in this kit. The BAL sample data were corrected for the dilution factor since 25 µL was used versus 10 µL of the standards that were used to calculate the quantitative calibration curve. Principal component analysis of log normalized counts or concentrations were performed in R (version 4.0.2) (Team, 2021) using the prcomp() function and visualized with ggplot2 (Wickham, 2016) using ggfortify (Tang & Horikoshi, 2016). Supplemental table of sample metadata compiled with sjPlot (Lüdecke, 2021) in R.

### BIOLOG Phenotyping assay

Two mL of LB overnight cultures grown at 37°C on a roller drum were washed twice with M63 salts with no carbon source by repeated centrifugation (10 min, 13K RPM) and resuspension into fresh medium. The washed cultures were normalized to an OD_600 nm_ = 0.05 in 25 mL of fresh M63 salts base and 100 µL was used to resuspend dehydrated carbon sources on the bottom of PM1 and PM2 BIOLOG phenotype plates by repeated pipetting. Cells and resuspended carbon were transferred to a sterile flat bottom, black-walled 96 well plate and incubated at 37°C, static. Every hour OD_600 nm_ was monitored in a plate reader for 24 h. Endpoint (24h) data is reported. Data visualization and statistical analysis performed in GraphPad Prism 9 (version 9.2.0).

### Contact for reagents, data, resource sharing and code availability statement

All data necessary for evaluation of the manuscript conclusions are available within the main text or supplementary materials. Further information and requests for resources and reagents should be directed to and will be fulfilled by the corresponding author. No custom code was used.

## Acknowledgements

Research reported in this publication was supported by grants from the Cystic Fibrosis Foundation HOGAN19G0 (D.A.H.), ASHARE20P0 (A.A.) and STANTO19R0 (D.S.), and the National Institutes of Health (NIH) through T32AI007519 (D.L.M.), R01HL122372 (A.A.) and P20 GM130454-02 (D.S.). Additional core facility support came from the NIH NIGMS P20GM113132 (BioMT) and NIDDK P30-DK117469 (Dartmouth Cystic Fibrosis Research Center) and STANTO19R0 from the Cystic Fibrosis Foundation. Plasmid sequencing was carried out at Geisel School of Medicine Genomics Shared Resource, which was established by equipment grants from the NIH and NSF and is supported in part by a Cancer Center Core Grant (P30CA023108) from the NIH National Cancer Institute. We would like to acknowledge Amy Conaway for assistance in the kinase evolution screen, Dr. Georgia Doing for LasR^-^ colony enumeration in key experiments, and Dr. Nicholas Jacobs for constructive and thoughtful feedback on the written manuscript. The content is solely the responsibility of the authors and does not necessarily represent the official views of the NIH. The funders had no role in study design, data collection and analysis, decision to publish, or preparation of the manuscript.

---

**Supplemental Figure 1.**
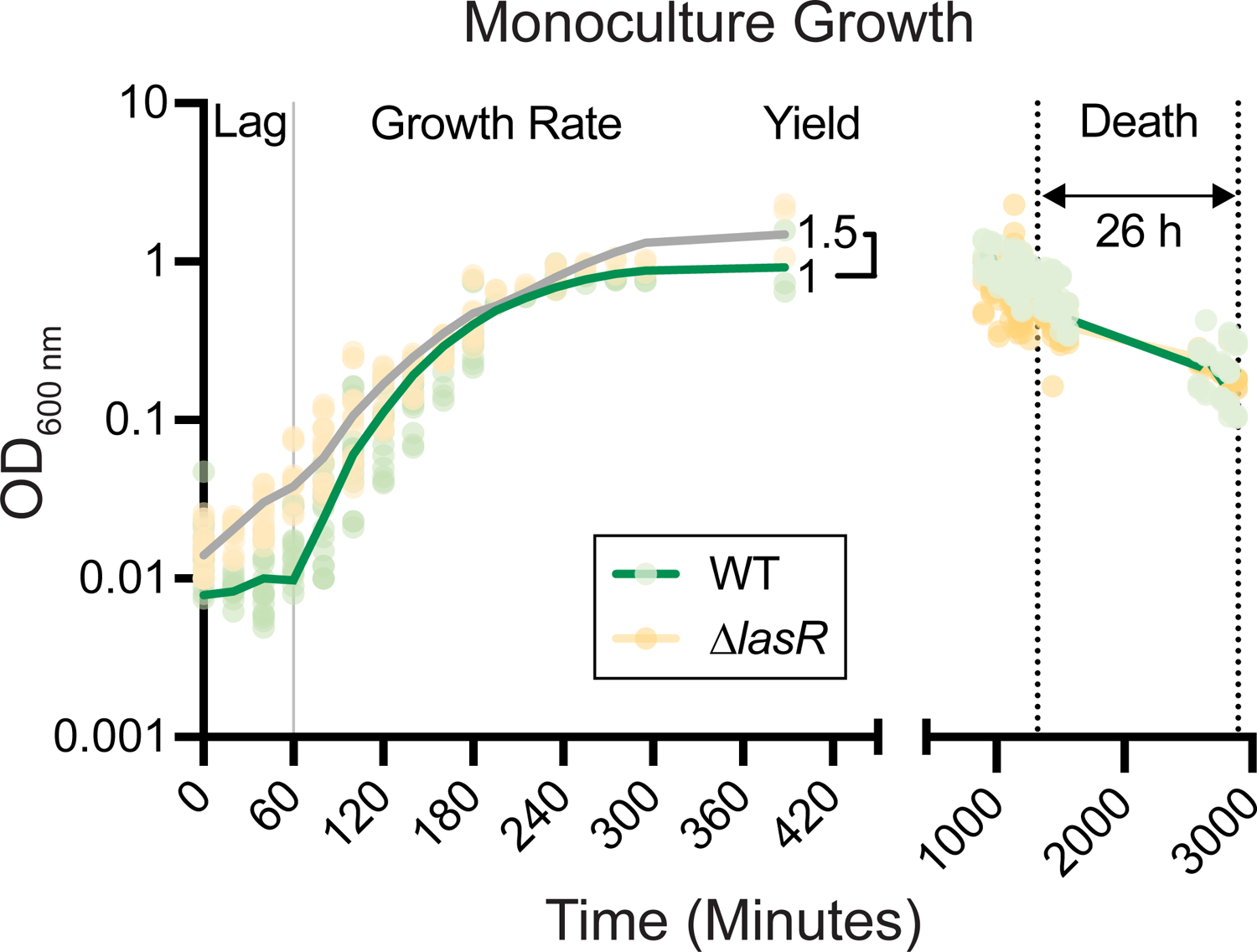
Experimentally determined growth parameters of PA14 wild type and Δ*lasR* monocultures in LB. Density measurements of strains PA14 wild type (WT, green) and Δ*lasR* (beige, grey) monocultures over time. Lag, growth rate, yield, duration of death, and death rate were determined for use in the mathematical model (Fig. 1A).

**Supplemental Figure 2.**
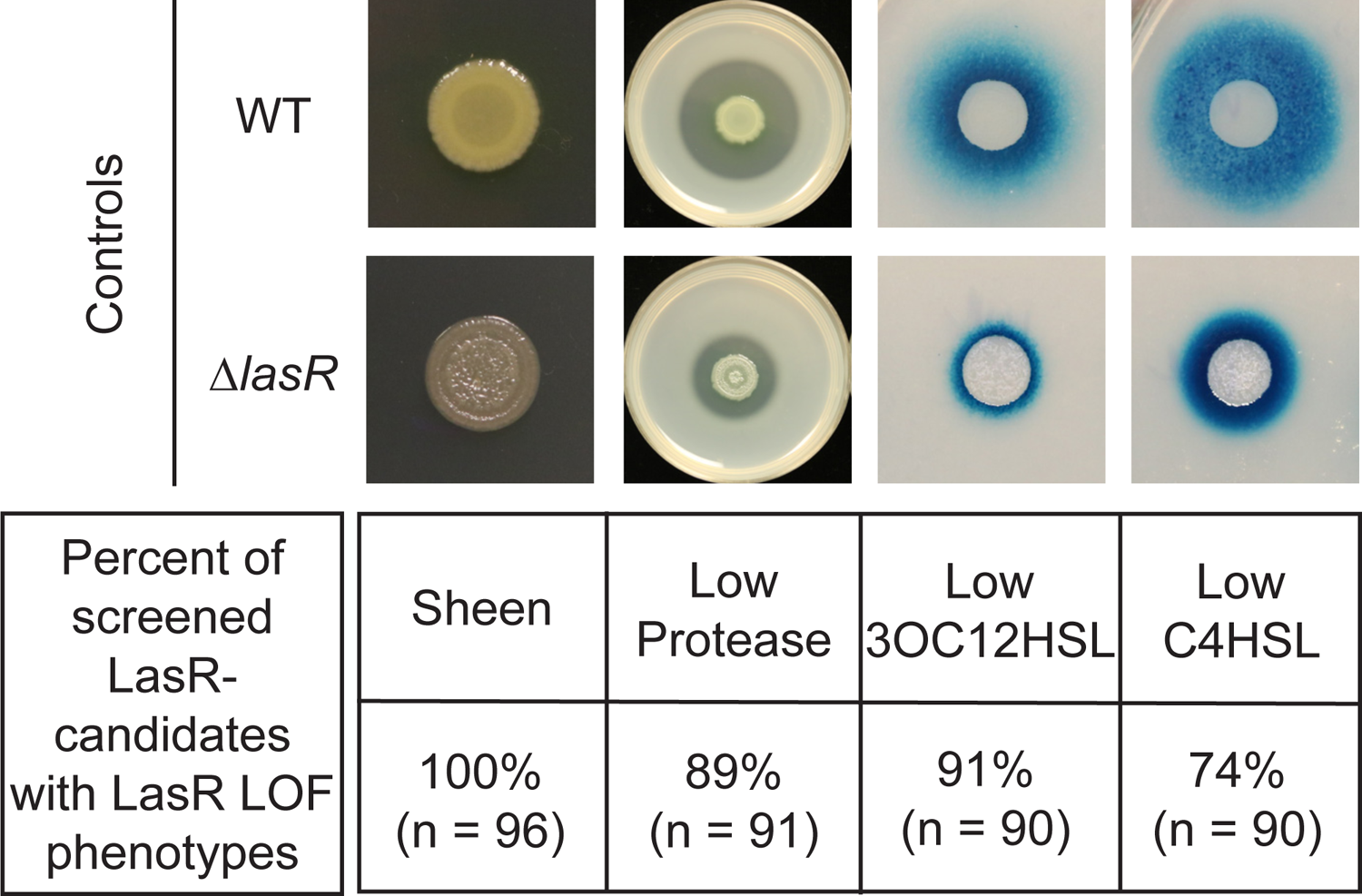
Phenotype analysis of sheen LasR-candidates isolated from the evolution experiments in LB. LasR loss-of-function (LOF) candidates picked on basis of sheen colony morphology from evolution were screened for phenotypes associated with LasR-strains including low protease activity on milk plates and low levels of acyl homoserine lactone production as measured by Δ*lasI*Δ*rhlI* bioreporters responsive to LasR-regulated autoinducer 3-oxo-C12-homoserine lactone (3OC12HSL) and RhlR-regulated autoinducer N-butyryl-L-homoserine lactone (C4HSL).

**Supplemental Figure 3.**
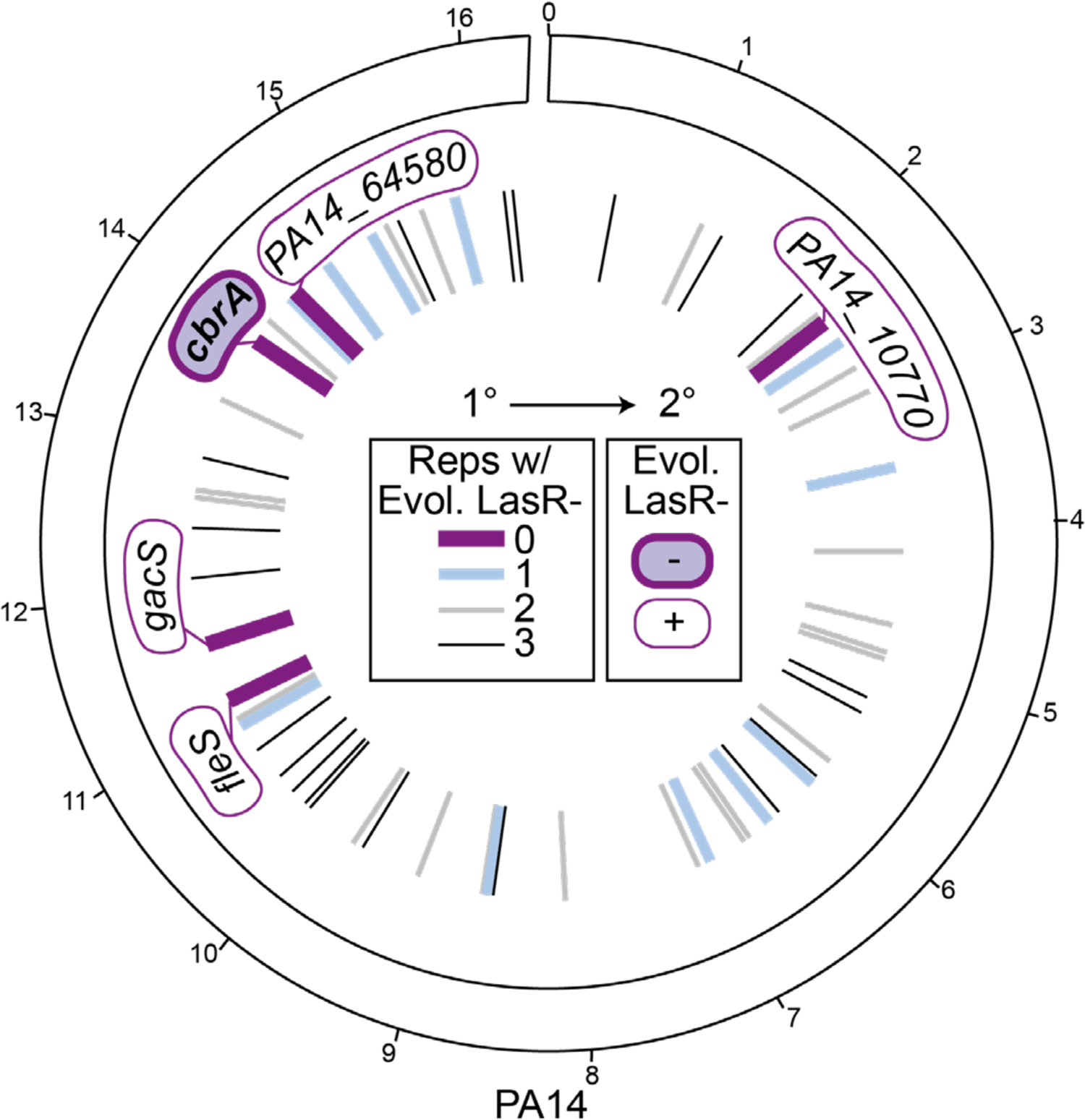
Screen reveals specific requirement of CbrA, and not other sensor kinases encoded in the PA14 genome for LasR-strain selection. Circos plot of the PA14 genome, with the genomic location of the genes deleted to create the kinase mutant collection indicated by lines in the inner circle (and noted in Table S2); outer ticks indicate 4 x 10^5^ bp genome increments. The primary screen (1°) was performed in a 96-well plate format in LB with each strain in triplicate. Deletion backgrounds that had zero, one, two, or three replicates containing LasR-phenotypes at the time of plating (i.e. when all wild-type controls had > 50% colonies with LasR-phenotypes) are represented as lines colored purple, blue, or grey respectively with decreasing line thickness. Deletion mutants that were found to have zero colonies with LasR-phenotypes (thick dark purple lines, gene names indicated) were secondarily screened in 5 mL LB evolution assays (Fig. S4A); only Δ*cbrA* (filled in purple) was negative (-) for LasR-phenotypes in the secondary screen (2°).

**Supplementary Figure 4.**
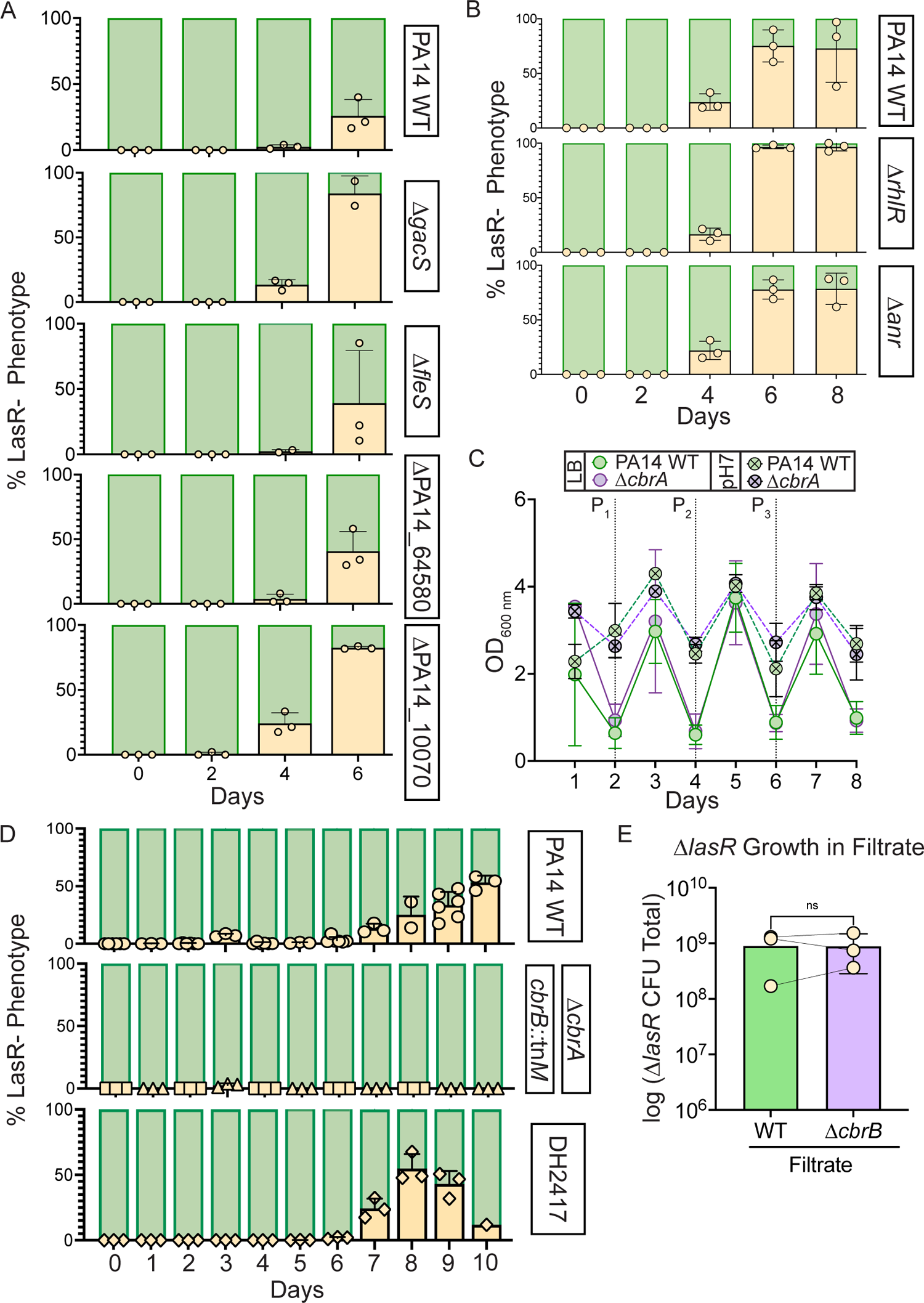
LasR-strains evolve in all tested strain backgrounds except for those deficient in *cbrAB,* and this is independent of cellular density, lysis, and filtrate toxicity. A. Kinase deletion mutant backgrounds that did not evolve LasR-phenotypes in any replicate of a 96-well evolution (Fig. S3) were secondarily screened for the appearance of LasR-colonies in 5 mL cultures. The colony phenotypes were quantified over time with percentage of LasR-phenotypes indicated (beige). B. The percent of colonies with LasR-phenotypes observed for LB evolution experiments initiated with strains PA14 Δ*rhlR* or Δ*anr* relative to wild type. C. 0ptical density for PA14 wild type (WT, green) or Δ*cbrA* (purple) cultures over course of evolution in LB (circles, n 2: 9) or LB buffered to pH 7 with HEPES (circle with “x”, n 2: 3). Points represent the average and error bars, standard deviation. Statistical significance between WT and Δ*cbrA* determined by two-way AN0VA with Sidak’s multiple comparisons test for LB and buffered LB datasets separately. For LB: Day 1, p < 0.0001. For buffered LB: Day 1, p = 0.023 and Day 6, p < 0.009. All other comparisons are non-significant. D. Percentage of colonies with LasR-phenotypes over the course of evolution in buffered LB for PA14 WT (circles), Δ*cbrA* (triangles) or *cbrB*::tn*M* (squares), or DH2417 LasR+ clinical isolate (diamonds) (n 2: 3). E. Density (i.e. total colony forming unit counts) of PA14 *!ilasR* after 24 h of growth in filtrate from saturated PA14 wild type or Δ*cbrB* cultures. ns, not significant as determined by Student’s t-test.

**Supplemental Figure 5.**
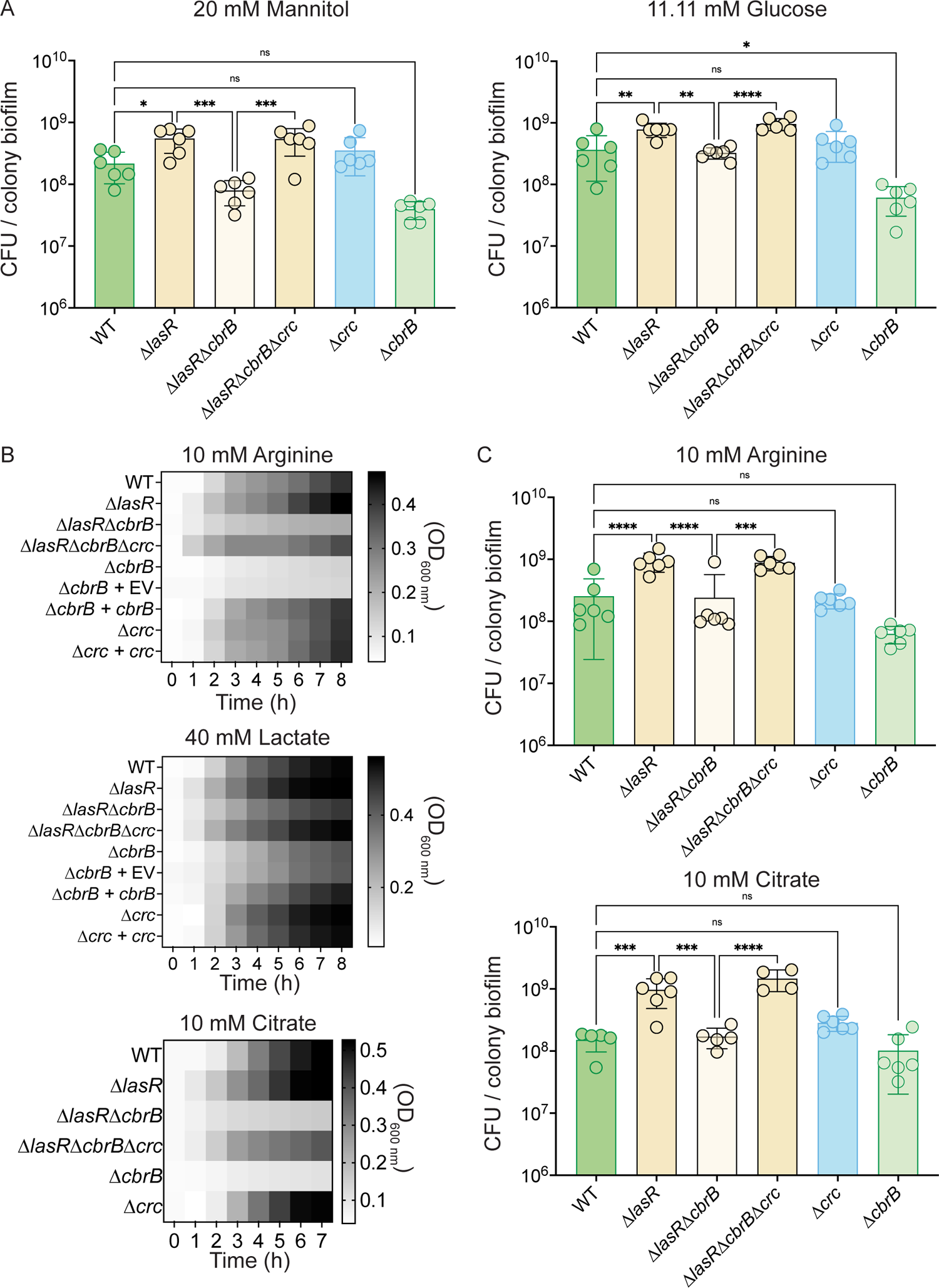
*AfasR* strains have CbrB-dependent growth advantages that can be restored via loss of *crc*. A. Colony biofilm CFUs were enumerated for strains PA14 WT, Δ*lasR,* Δ*lasR*Δ*cbrB,* Δ*lasR*Δ*cbrB*Δ*crc,* Δ*crc,* and Δ*cbrB* after 16 h on M63 minimal medium containing 20 mM mannitol or 0.2% glucose, which have been well studied in the context of carbon catabolite repression (i.e. the CbrAB pathway). B. Heatmap representation of planktonic growth on different carbon sources: 10 mM arginine, 40 mM lactic acid, and 10 mM citrate. Heatmaps show the average growth (OD _600 nm_) across three independent experiments with three replicates per day. C. Colony counts of resuspended colony biofilms grown on agar plates containing 10 mM Arginine or 10 mM citrate as sole carbon sources. Bottom of y-axis set to starting inoculum density. P-values: ns, not significant; *, p < 0.05; **, p <; ***, p < 0.0007; ****, p < 0.0001 as determined by ordinary one-way AN0VA with Sidak’s multiple comparisons test. Each data point of colony count (CFU) was collected on a separate day (n = 6).

**Supplemental Figure 6.**
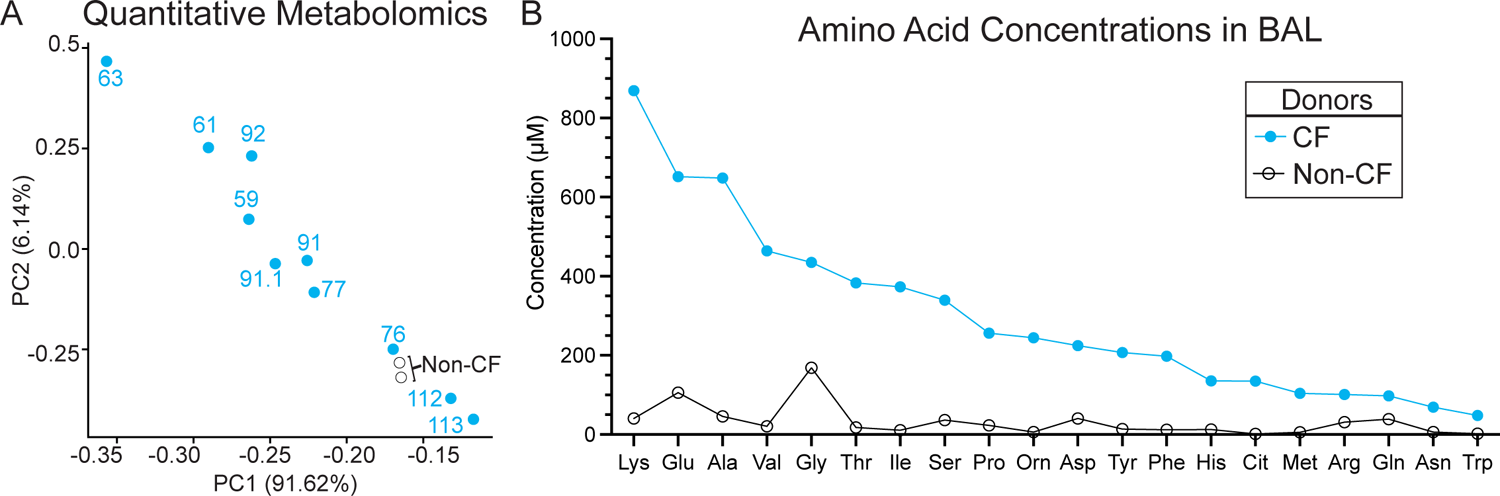
Quantitative amino acid analysis. A. The first two components (PC1 and PC2) of a principal component analysis of log normalized amino acid concentrations measured in bronchoalveolar lavage (BAL) luid collected from cystic fibrosis (CF, blue filled circles) and non-cystic fibrosis (Non-CF, black open circles) donors by the Biocrates AbsolutelDQ p180 Kit explain 91.62% and 6.14% of the variation in the data, respectively. As with the relative metabolite counts measured by LC/MS used in Fig. 4A, PC1 of the amino concentrations measured by the Biocrates AbsolutelDQ p180 Kit separates BAL samples by the respective percent forced expiratory volume in 1 sec (%FEV_1_) (overlayed text). B. Average amino acid concentrations (µM) measured from BAL samples collected from cystic fibrosis (CF, blue) and non-cystic fibrosis (Non-CF, black) donors. See Supplemental Table 5 for data by sample.

